# Human spastin functions as an ATPase-independent microtubule nucleator via ring-stack formation

**DOI:** 10.1101/2025.11.28.691112

**Authors:** Hanjin Liu, Nao Matsumoto, Toshiyuki Kaneshiro, Takaaki Kato, Hideki Shigematsu, Satoshi Kikkawa, Eriko Nitta, Tsuyoshi Imasaki, Ryo Nitta

## Abstract

Microtubule severing enzyme spastin plays pivotal roles in cytokinesis and neuronal outgrowth, and its mutations cause hereditary spastic paraplegia (HSP). Here we show that, at physiological tubulin concentrations, human spastin behaves predominantly as an ATPase-independent microtubule nucleator rather than a severase. Biochemical and structural analyses revealed that spastin assembles tubulin into stacked-ring intermediates via strategically positioned microtubule-binding domains, thereby generating nucleation-competent sites for polymerization. Tubulin subunits within these spastin-induced rings adopt a straight, microtubule-like interface, in marked contrast to the twisted tubulin spirals characteristic of depolymerizing microtubule ends. Our results redefine the mechanistic landscape of multifaceted spastin functions and provide insights that may inform the pathological basis of HSP and guide future therapeutic strategies.

## Main Text

The microtubule cytoskeleton is an indispensable component of eukaryotic cells, supporting cell shape and driving dynamic processes such as chromosome segregation and cell locomotion (*1*). These diverse functions emerge from continuous reorganization of microtubules through the addition and removal of αβ-tubulin subunits, a process tightly regulated by numerous microtubule-associated proteins (MAPs) (*2*). MAPs mediate finely controlled growth and shrinkage at microtubule ends (*3*, *4*), redistribution of microtubules (*5*, *6*), and de novo nucleation of microtubules from tubulin subunits (*7*, *8*). Among MAPs, microtubule severing enzymes, spastin, katanin, and fidgetin, are AAA+ hexameric ATPases that cut microtubules within the lattice, most likely by extracting the tubulin C-terminal tail through the central pore of the hexamer (*9–16*). This activity promotes controlled microtubule turnover in processes such as mitotic spindle disassembly and nuclear envelope resealing (*17*, *18*). It has also been harnessed as a tool for manipulating microtubules in living cells (*19*, *20*). Spastin, in particular, is necessary for neuronal outgrowth (*21–23*), abscission (*24*, *25*), and nuclear envelope resealing during mitosis (*17*). Importantly, mutations in the spastin gene represent the most common cause of hereditary spastic paraplegia (HSP) (*26*).

Dysregulation of spastin causes significant impacts on cells, but seemingly produces opposing outcomes at the level of microtubule network. Overexpression depletes microtubules in cultured human cells (*10*), whereas spastin knockout leads to denser microtubule arrays in mouse neurons (*27*). In contrast, spastin downregulation disrupts microtubule network formation in zebrafish (*28*) and reduces the population of stable microtubules in mice (*29*). Thus, spastin is not merely a microtubule breaker but can also act as a network builder, with its dual roles depending on the precise cellular context and concentration.

In vitro reconstitution studies suggested mechanisms that may reconcile this paradox. Mild microtubule severing can increase the number of microtubule fragments that serve as seeds for regrowth (*30*). More recently, studies using *Drosophila* spastin demonstrated that partially severed microtubules can undergo lattice repair via incorporation of free tubulin, thereby creating rescue sites that protect microtubules from complete depolymerization (*31*)– a phenomenon broadly referred to as lattice repair (*32*, *33*). In addition, *Drosophila* spastin has been shown to reduce the shrinkage rate and increase the rescue frequency of depolymerizing microtubules in an ATPase-independent manner (*34*). These reports establish the concept of a “severing-dependent microtubule amplification mechanism,” whereby spastin activity can increase overall microtubule mass.

However, the feasibility of this mechanism depends critically on the spatial and temporal precision of severing, as uncontrolled activity inevitably leads to catastrophic microtubule loss. Although several regulatory factors—including microtubule polyglutamylation (*35*), spastin phosphorylation (*36*), and tau-mediated microtubule protection (*37*)—have been identified in modulating severing activity, how cells balance the opposing outcomes of spastin action, degeneration versus mass amplification, remains unanswered at the molecular level. Here we delineate a previously unrecognized, severing- and template-independent pathway for microtubule amplification in which human spastin hexamers organize tubulin into stacked rings and spirals that serve as nucleation seeds. High-resolution structural analyses further identify that symmetry-mismatched ring stacking and protofilament untwisting are key determinants of this spastin-driven nucleation mechanism.

### Free tubulin inhibits severing activity of human spastin via tubulin ring formation

The function of microtubule severing by spastin has been extensively studied using constructs from *Drosophila* and human (*10*, *13*, *14*, *38*, *39*). However, multiple sequence alignments reveal a clear divergence between Protostomia and Deuterostomia, suggesting evolutionary and functional divergence (Fig. 1A; fig. S1). While the core AAA ATPase domain is highly conserved, the microtubule-binding domain (MTBD)—which is necessary and sufficient for microtubule binding (*39*, *40*), and strongly influences the cellular behavior of severing proteins (*41*) (Fig. 1B)—differs markedly between human and *Drosophila*. Notably, the human MTBD contains highly positively charged residues as well as the threonine-rich region (Fig. 1A).

**Fig. 1.**
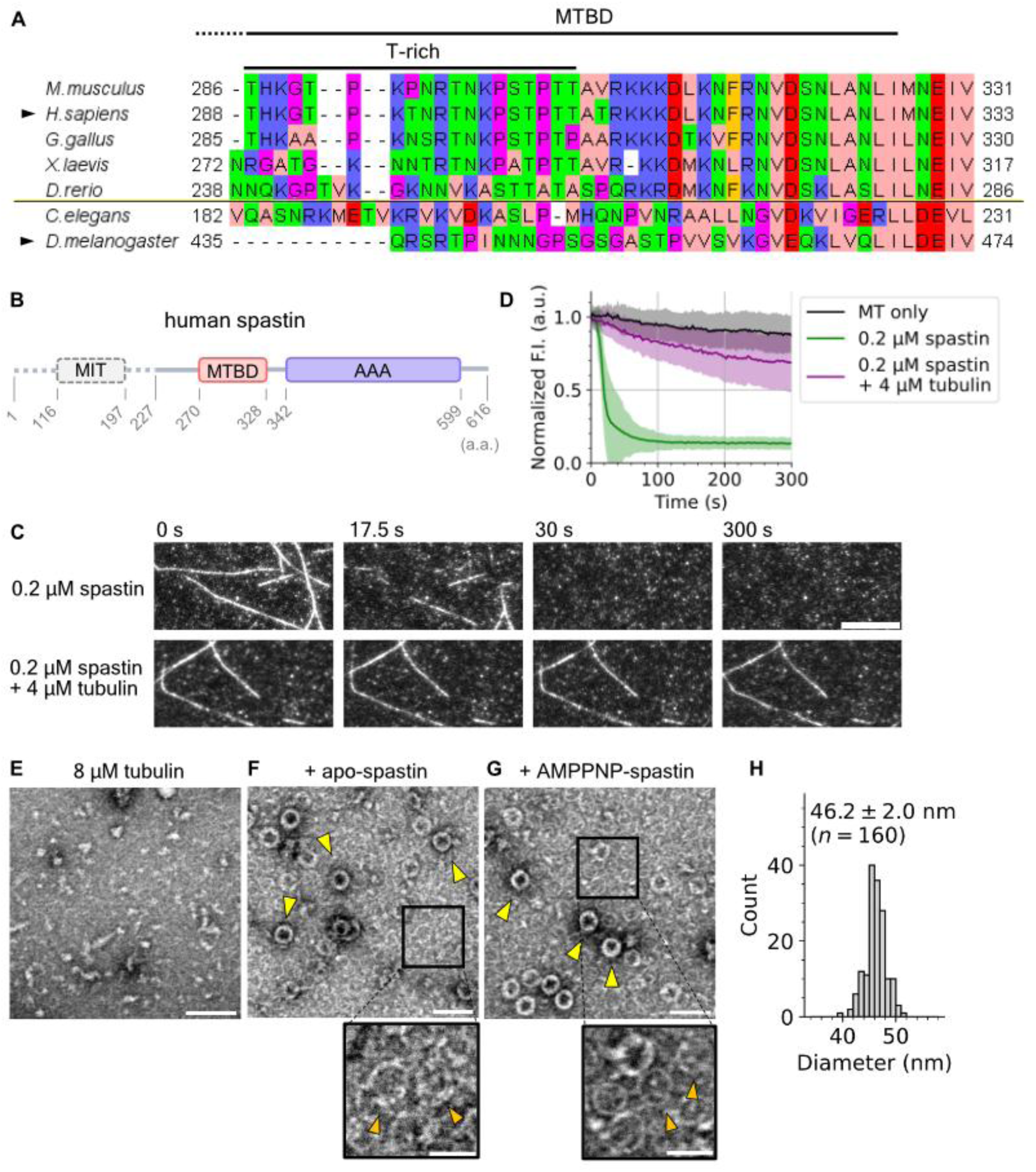
Microtubule severing by human spastin was inhibited by free tubulin with the formation of tubulin rings. (**A**) Multiple sequence alignment of the sequence around the MTBD of human spastin. Horizontal line splits Protostomia (lower) and Deuterostomia (upper). Alignment and visualization were performed by JalView (v2.11.4.1). (**B**) Domain structure of human spastin. The first 227 residues are not included in the spastin construct used in this study, as this region is irrelevant to the microtubule severing activity(*39*). (**C**) Representative time-lapse images of spastin severing assay at 28°C. Scale bar, 10 µm. (**D**) Quantification of severing activity using the fluorescence intensity (F. I.) of microtubules normalized by the initial value. (**E–G**) Negative-stain EM micrographs of spastin-dependent tubulin ring formation at 37°C. Boxed regions are zoomed in the lower panels. Arrowheads point at thick tubulin rings (yellow) and thin tubulin rings (orange). Scale bar, 100 nm (upper) and 50 nm (lower). (H) Histogram of the diameter of tubulin rings formed by apo-spastin. 160 rings in eight micrographs were analyzed.

Based on these comparisons, we hypothesized that the tubulin-binding properties of spastin vary across species. In *in vitro* reconstitution assays, free tubulin slows the microtubule severing rate of *Drosophila* spastin (DmSpastin) by partially suppressing microtubule-binding (*31*, *34*), but differences in the MTBD may cause human spastin to behave differently. To test this, we quantified the severing activity of human spastin in the absence and presence of free tubulin and evaluated the extent to which soluble tubulin in solution inhibits severing activity (Fig. 1B). Severing of fluorescently labeled, taxol-stabilized microtubule was observed using total internal reflection fluorescence microscopy (TIRFM), as previously applied to katanin (*42*), and was quantified by decreases in fluorescence intensity of microtubules.

In the absence of free tubulin, 0.2 µM human spastin—half the concentration used in White et al. (*39*) —effectively severed microtubules, leading to their complete disappearance within ∼20 s (Fig. 1C). By contrast, severing activity was strongly suppressed in the presence of 4 µM free tubulin over 5-min (Fig. 1, C and D). This behavior differs markedly from DmSpastin, which severs microtubules even at 12 µM tubulin (*31*, *34*). The normalized fluorescence intensity decreased more rapidly at 0.2 µM spastin with 4 µM tubulin than in microtubule-only controls, suggesting lattice damage from partial severing followed by repair (*31*). Indeed, spastin-dependent tubulin incorporation into the microtubule lattice was observed by TIRFM-based repair assay (fig. S2).

We assumed that induction of lattice damage is a conserved feature of microtubule severing enzymes, and thus expected human spastin to behave similarly. However, the inhibitory effect of free tubulin appears substantially stronger for human spastin than for DmSpastin, such that the rate of lattice repair matches the rate of severing. Since the balance between tubulin-binding and microtubule-binding determines the extent of inhibition by free tubulin, our results suggest that human spastin possesses additional tubulin-interaction modes not detected in assays using DmSpastin.

To directly visualize the interaction between tubulin and human spastin (hereafter, spastin), we co-incubated these proteins at 37°C and observed the mixtures by negative-stain electron microscopy (EM). Intriguingly, across a range of spastin concentrations, we observed multiple ring-shaped structures in a spastin concentration-dependent manner (Fig. 1, E and F; fig. S3A). Rings formed regardless of the spastin’s nucleotide states, including apo, AMPPNP, ADP-vanadate (ADP-Vi), and ADP (Fig. 1, F and G; fig S3, B and C), indicating that ring formation is independent of the ATPase activity of spastin.

Each field of view contains low-contrast “thin rings” and high-contrast “thick rings”, suggesting that these rings can enlarge either by overlapping or by spirally rolling to form double or triple rings, which is often observed in polymerizing or depolymerizing microtubules (*8*, *43*) (Fig. 1, F and G). The mean diameter of the thick rings was 46.2 ± 2.0 nm (Fig. 1H), corresponding to ∼20°/dimer curvature—well within the range of longitudinal tubulin–tubulin interactions, such as those at elongating or shortening protofilament tips (10 – 30°/dimer) (*44*).

Strikingly, microtubules were also found in the grids containing spastin, even though the tubulin concentration was below the critical concentration for microtubule polymerization, suggesting that spastin promotes microtubule formation (fig. S3D). Consistent with this, pelleting assay at 4°C—conditions under which tubulin does not assemble into microtubules—showed spastin-dependent formation of tubulin rings or larger assemblies (fig. S3E). Together, these observations demonstrate that human spastin assembles tubulin into ring complexes in solution. This interaction likely underlies the pronounced inhibitory effect of free tubulin on spastin’s severing activity.

### Spastin nucleates microtubules independently of ATP hydrolysis

Our observation that free tubulin potently inhibits microtubule severing by human spastin argues against severing-dependent amplification in human cells (*31*, *34*). We therefore examined an alternative mechanism centered on the ring structures. Although ring-shaped tubulin oligomers have been linked to microtubule depolymerization (*45*, *46*), our TIRFM-based depolymerization assay showed that human spastin suppresses depolymerization (fig. S4), consistent with findings for DmSpastin (*34*). This indicates that the rings we observe are not the depolymerization products. Notably, tubulin rings have long been reported (*43*, *47*) and frequently arise under nucleation-prone conditions, including co-incubation with taxol (*48*), CLIP-170 (*49*), doublecortin (*50*), CAMSAP2 (*8*), and GAS2 (*51*). Together with our observation that spastin promotes formation of both tubulin rings and microtubules (Fig. 1, E–G; fig. S3), these findings led us to hypothesize that human spastin acts as a microtubule nucleator through ring formation.

We first tested this hypothesis using turbidity assays that monitored the increase of light scattering at 350 nm (*52*) with 15 µM tubulin, a concentration at which spontaneous nucleation is rare (*8*, *53*, *54*). Tubulin alone produced only a weak signal after 30 min (Fig. 2A), whereas taxol induced rapid polymerization (fig. S5). Strikingly, 1 µM spastin induced microtubule formation in the absence of ATP and more efficiency in its presence (Fig. 2A). Negative stain EM of post-assay samples confirmed bona fide microtubules (Fig. 2B). The end-point absorbance of spastin-treated samples exceeded that of taxol polymerized samples, consistent with spastin-induced bundling (*55*), because light scattering is sensitive to the diameter. Nucleation was also detectable at 0.5 µM spastin, albeit with less efficiency in the absence of ATP, similar to the behavior observed at 1 µM.

**Fig. 2.**
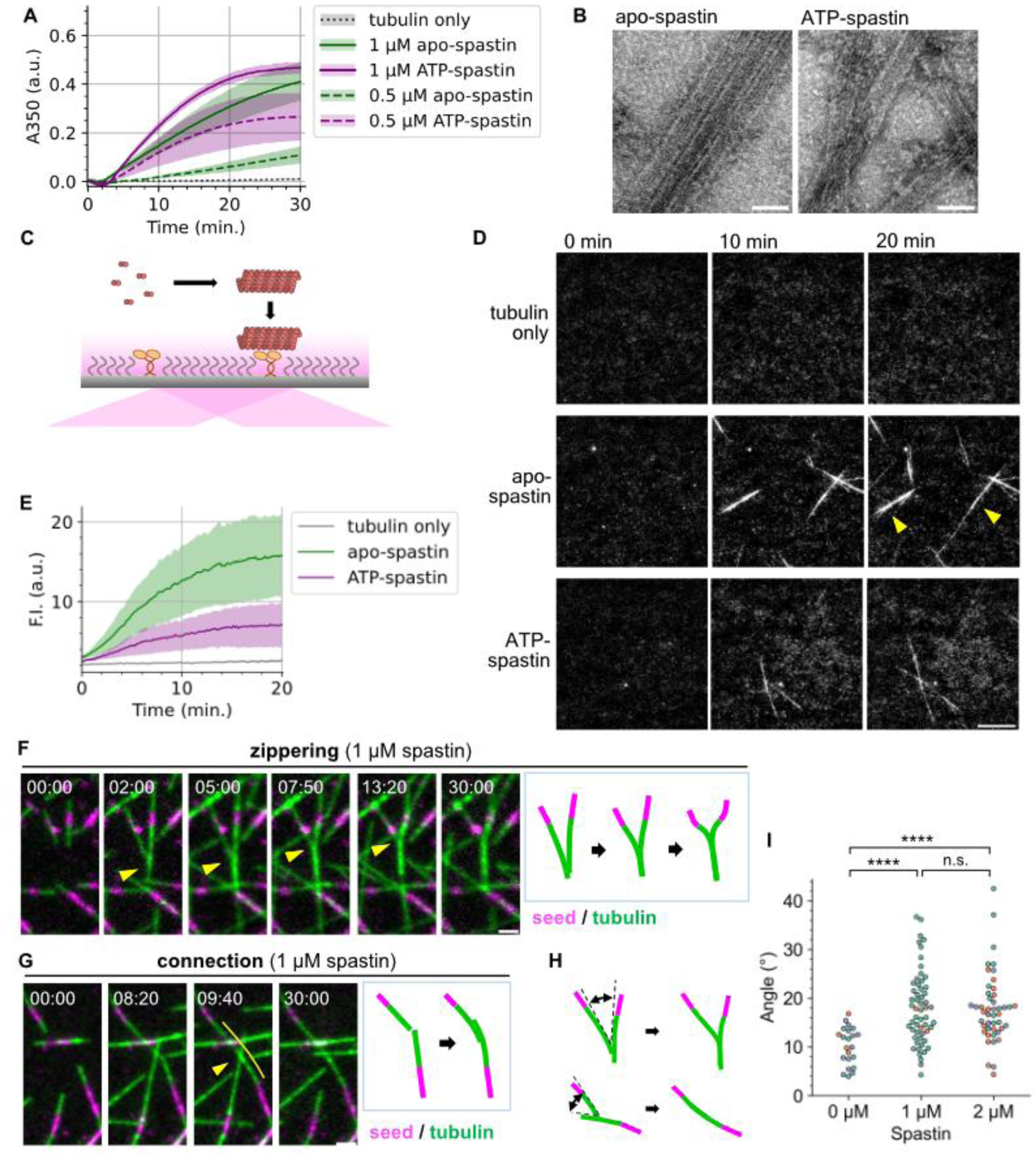
The nucleation and bundling activity of human spastin. (**A**) Turbidity assay showing the nucleation activity of spastin. 15 μM tubulin was co-incubated with 0.5 µM (dotted curves) or 1 µM (solid curves) spastin, without (green) or with (purple) 1 mM ATP. Line and shaded area indicate mean ± SE (*n* = 3). (**B**) Negative-stain micrographs of the samples after turbidity assay of tubulin with 1 µM spastin. Scale bar, 50 nm. (**C**) Schematic illustration of TIRF-based nucleation assay. KIF5C only binds to polymerized tubulin, thus the increase in the microtubule mass can be monitored by the total fluorescence intensity. (**D**) Representative fluorescence images of the TIRF-based nucleation assay at 32°C. Arrowheads indicate the bundled microtubules. Scale bar, 10 µm. (**E**) Quantification of the increase of fluorescence intensity of microtubules. Solid line and shaded area indicate mean ± SE (*n* = 3). (**F**) Representative image series of the zippering event. Arrowheads indicate the branching point. Scale bar, 2 µm. (**G**) Representative image series of the connection event. Arrowhead indicates the point where bundling occurred. Scale bar, 5 µm. (**H**) Schematic illustration showing the angles between microtubules just before zippering or connection. (**I**) Plot of the angles indicated in (H). *n* = 22, 56 and 46 events for 0, 1 and 2 µM spastin, respectively. Color represents each replicate of three independent experiments. n.s.: not significant, ****: *p* < 0.0001 (Steel-Dwass test).

To directly visualize nucleation, we monitored spastin-driven assembly using TIRFM (Fig. 2C). With 15 µM tubulin, 1 µM spastin robustly promoted microtubule formation even in the absence of ATP (Fig. 2D), consistent with turbidity data. No severing events were detected, most likely because free tubulin suppresses the severing activity of spastin even when ATP is present, indicating that severing is not required for microtubule mass amplification under these conditions. Quantified fluorescence time courses reproduced the overall trends of the turbidity assay (Fig. 2E).

We also observed bundling of microtubules in this nucleation assay (Fig. 2D) and TIRFM-based bundling assay (fig. S6, A and B), as suggested from the high absorbance in the turbidity data (Fig. 2A). The only notable discrepancy was an inversion of apparent apo- versus ATP-spastin efficiencies, which we attribute to assay-specific biases: turbidity is enhanced by bundling, whereas in TIRFM, ATP-dependent differences in microtubule affinity may reduce apparent nucleation if spastin occupancy competes with kinesin-mediated surface attachment. Together, these results demonstrate ATP-independent microtubule nucleation by spastin, with nucleation predominating over severing in our conditions.

### Spastin reorganizes the microtubule network by bundling

After nucleation, tubulin polymerizes on newly formed microtubule seeds. In turbidity assays, this elongation phase appears as a linear increase in microtubule mass until reaching a plateau (*48*, *56*) (Fig. 2A). Microtubule bundling observed in TIRFM (Fig. 2D) suggests that spastin nucleates highly ordered microtubule arrays instead of a collection of randomly oriented microtubules. To test this, we conducted microtubule dynamics assays specifically during the polymerization phase. Stabilized microtubule seeds were immobilized on the glass surface, and elongation trajectories from seeds were monitored in the presence of spastin and ATP.

At 1 – 2 µM spastin, severing events were rare during the 30-min assays. When severing did occur, the resulting fragments were rapidly “patched,” likely through re-bundling (fig. S6C). We also observed pulling events in which a depolymerizing microtubule reoriented a neighboring filament (fig. S6D), as well as diffusion of tubulin puncta along existing microtubules, consistent with the diffusive binding mode of spastin (*40*) coupled to tethering of tubulin ring stacks (fig. S6E).

Bundling was prominent in the presence of spastin. When two microtubules elongated in near-parallel orientations, they underwent “zippering” (Fig. 2F). When they elongated in near-antiparallel orientations, “connection” events occurred, which can also arise without spastin (Fig. 2F; fig. S6F). Similar behaviors have been reported for an artificial tetrameric microtubule binder (*57*) and for tau (*58*), suggesting that multivalent microtubule interactions drive these events.

To quantify spastin’s contribution, we measured the angle between microtubules immediately before zippering or connection (Fig. 2H). To avoid overestimating angles, freely diffusing, newly nucleated microtubules were excluded (fig. S6G). Compared with control, the addition of 1 or 2 µM spastin almost doubled the maximum angle at which microtubules could bundle (Fig. 2I). These results indicate that, once sufficient nuclei are present, spastin functions as a microtubule network remodeler that promotes formation of parallel microtubule bundles.

### ATPase and severing activities are dispensable for ring formation and nucleation

As nucleation is a prominent activity of spastin in the presence of free tubulin, we next sought to define its molecular mechanism. Because nucleation occurred without ATP, we hypothesized that spastin-tubulin interactions alone—rather than ATPase or severing activities—are sufficient for ring formation and nucleation. We therefore examined two severing-deficient mutants, E442Q and C448Y, of which only C448Y retains ATPase activity (*10*, *39*, *59*). Both mutants, purified to homogeneity (fig. S7A), failed to sever microtubules even at 2 µM (fig. S7, B and C). Negative stain EM revealed tubulin rings formed by both mutants with or without ATP, although apo-E442Q produced thinner rings (Fig. 3A). Turbidity assays confirmed nucleation activity for both mutants (Fig. 3B; fig. S8C), with distinct ATP dependence at 0.5 µM: E442Q was ATP-activated, similar to wild-type, whereas C448Y was ATP-deactivated. This behavior parallels the reduced mass of stable microtubules observed in C448Y-expressing mice (*29*). We therefore conclude that the ATP state of spastin modulates nucleation efficiency, but neither ATPase nor severing activity is required for tubulin ring formation or microtubule nucleation.

**Fig. 3.**
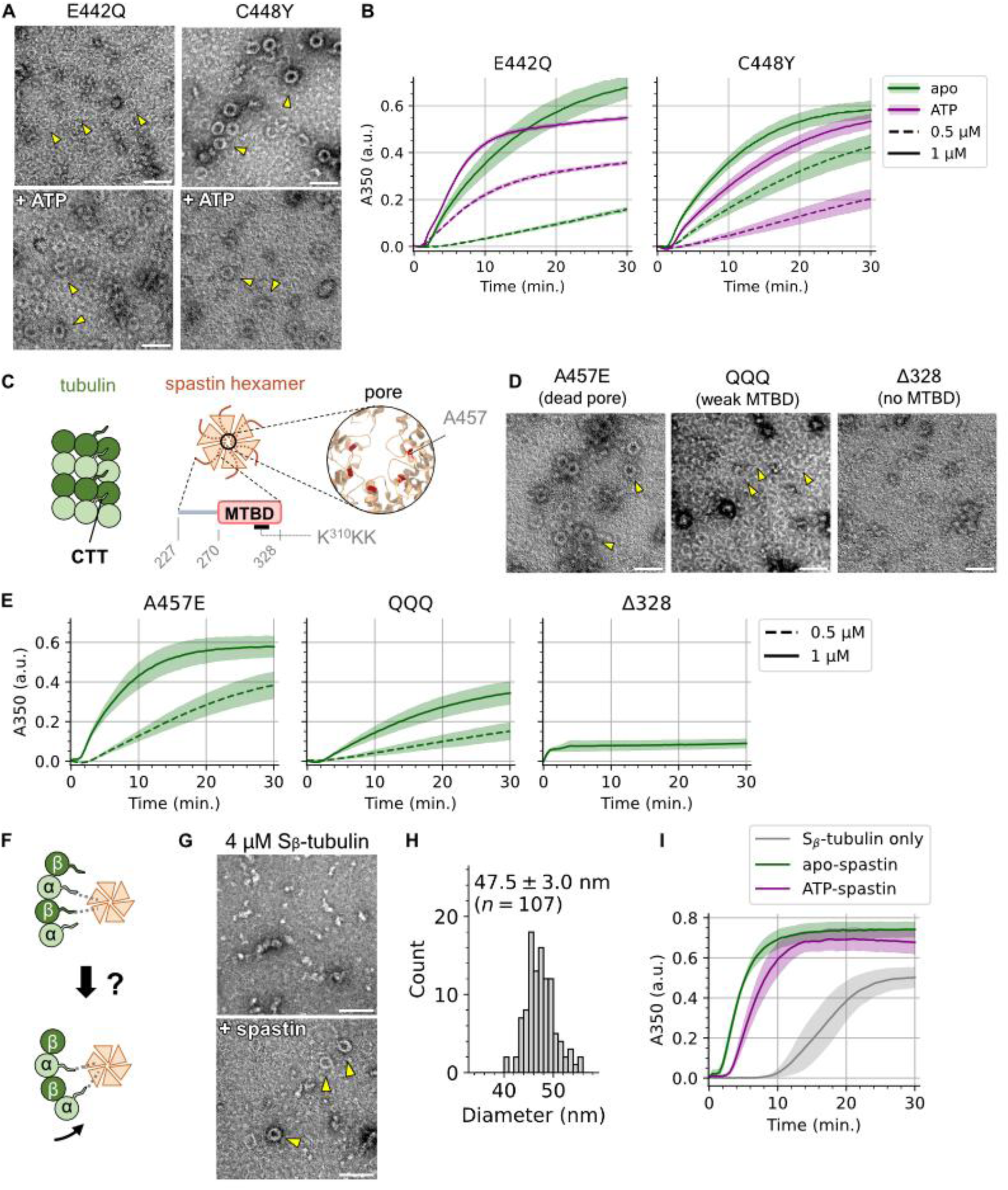
Identification of the spastin-tubulin interaction important for the ring formation. (A) Negative-stain EM micrographs of tubulin rings formed by severing deficient spastin mutants, E442Q and C448Y, at 37°C. Arrow heads point at the tubulin rings. Scale bar, 100 nm. (B) Results of turbidity assay using E442Q and C448Y. 15 µM tubulin was co-incubated with 0.5 µM (dotted curves) or 1 µM (solid curves) spastin mutants, without (green) or with (purple) 1 mM ATP. Line and shaded area indicate mean ± SE (0.5 µM apo-C448Y: *n* = 4; others: *n* = 3). (C) Schematic illustration of tubulin and spastin hexamer. Possible interactions are between tubulin CTT and spastin MTBD, and between tubulin CTT and the spastin pore. The K^310^KK residues shown here are mutated to glutamates in the QQQ mutant. (**D**) Negative-stain EM micrographs of tubulin rings formed at 37°C by spastin mutants whose interaction with tubulin is possibly blocked. Arrow heads point at the tubulin rings. Scale bar, 100 nm. (**E**) Results of turbidity assay using A457E, QQQ and Δ328. 15 µM tubulin was co-incubated with 0.5 µM (dotted curves) or 1 µM (solid curves) spastin mutants. Line and shaded area indicate mean ± SE (*n* = 3). (**F**) Schematic illustration of a possible effect of loss of the β-tubulin CTT. (**G**) Negative-stain micrographs showing the spastin-dependent S_β_-tubulin ring formation at 37°C. Arrowheads point at tubulin rings. Scale bar, 100 nm. (**H**) Histogram of the diameter of S_β_-tubulin rings. 107 rings in eight micrographs were analyzed. (**I**) Turbidity assay using S_β_-tubulin. Line and shaded area indicate mean ± SE (*n* = 3).

### MTBD–α-tubulin CTT interaction drive ring formation and nucleation

We next mapped the interaction responsible for ring formation and microtubule nucleation. Two spastin-tubulin interfaces have been described *in vitro*: the interaction between tubulin C-terminal tails (CTTs) and the spastin MTBD, and the interaction between tubulin CTTs and the spastin pore (Fig. 3C) (*39*). To dissect their respective contributions, we inhibited MTBD-CTT binding using the QQQ mutant, in which three consecutive lysine residues (K^310^KK) are replaced by glutamines (Q), or eliminated the MTBD entirely with the Δ328 truncation. To perturb pore-CTT contacts, we used A457E, which introduces electrostatic repulsion (*39*).

Severing assays showed that Δ328 and A457E were severing-deficient at 2 µM, whereas QQQ was only mildly impaired (fig S7, B and C). Negative stain EM demonstrated robust ring formation by A457E and QQQ, but not by Δ328 (Fig. 3D). Pelleting assays detected tubulin-spastin complexes for A457E and QQQ, but not Δ328 (fig. S8, A and B). Consistently, turbidity assays demonstrated nucleation activity for A457E and QQQ, but not for Δ328 (Fig. 3E; fig. S8C). Together, these data support a model in which spastin engages tubulin CTTs through its MTBD to assemble tubulin into rings, thereby promoting microtubule nucleation.

### A mechanistic model and β-CTT dispensability

Integrating the mutant data, we propose that spastin facilitates tubulin oligomerization into a microtubule nucleation seed through MTBD-CTT interactions, which are critical for nucleation efficiency (*60*). Given the size of tubulin monomer (55 kDa) and the spastin AAA domain monomer (32 kDa), we initially speculated that spastin might bridge adjacent β- and α-CTTs. If this were the case, removal of the β-CTT would be expected to prevent ring formation or reduce ring diameter (Fig. 3F).

To test this, we generated S_β_-tubulin (β-CTT–cleaved tubulin) by low-dose subtilisin treatment (*61*, *62*) (fig. S9A). Because S_β_-tubulin nucleates spontaneously at much lower concentration than intact tubulin, we performed negative stain EM and turbidity assay at reduced tubulin concentrations. Contrary to our expectation, S_β_-tubulin formed spastin-dependent rings (Fig. 3G) with diameters indistinguishable from those formed with intact tubulin (Fig. 3H; 47.5 ± 3.0 nm, while 46.2 ± 2.0 nm for the intact tubulin). Turbidity assays also confirmed that spastin promoted nucleation of S_β_-tubulin (Fig. 3I; fig. S9B). These results indicate that the β-CTT is dispensable for ring formation and nucleation. Moreover, because the β-CTT is essential for spastin-mediated severing, these findings further reinforce that severing is not required for nucleation (*35*, *38*).

### Cryo-ET reveals trivalent interactions of spastin with microtubules and tubulin ring stacks

Spastin functions as a hexamer, and tubulins assemble into rings and microtubules; however, the detailed binding modes and dispensability of the tubulin CTT remain unclear. To address these questions, we reconstructed the early microtubule nucleation intermediates formed with spastin and analyzed them by cryo-electron tomography (cryo-ET). Tomograms were denoised with Cryo-CARE (*63*), which is widely applied to *in vitro* reconstitution systems of microtubules and associated proteins (*64*, *65*). When 30 µM apo-spastin was co-incubated with 30 µM tubulin for 30 s at 37°C (Fig. 4A), numerous microtubules were already formed (Fig. 4B). Reconstructions revealed coexisting tubulin rings, sheets, and microtubules, together with densities corresponding to free or microtubule-bound spastin hexamers (Fig. 4, B–D; fig. S10). Consistent with the TIRFM observations (Fig. 2, F–I; fig. S6), bundled microtubules and zippering-like reorientation events were also captured in tomograms (fig. S10).

**Fig. 4.**
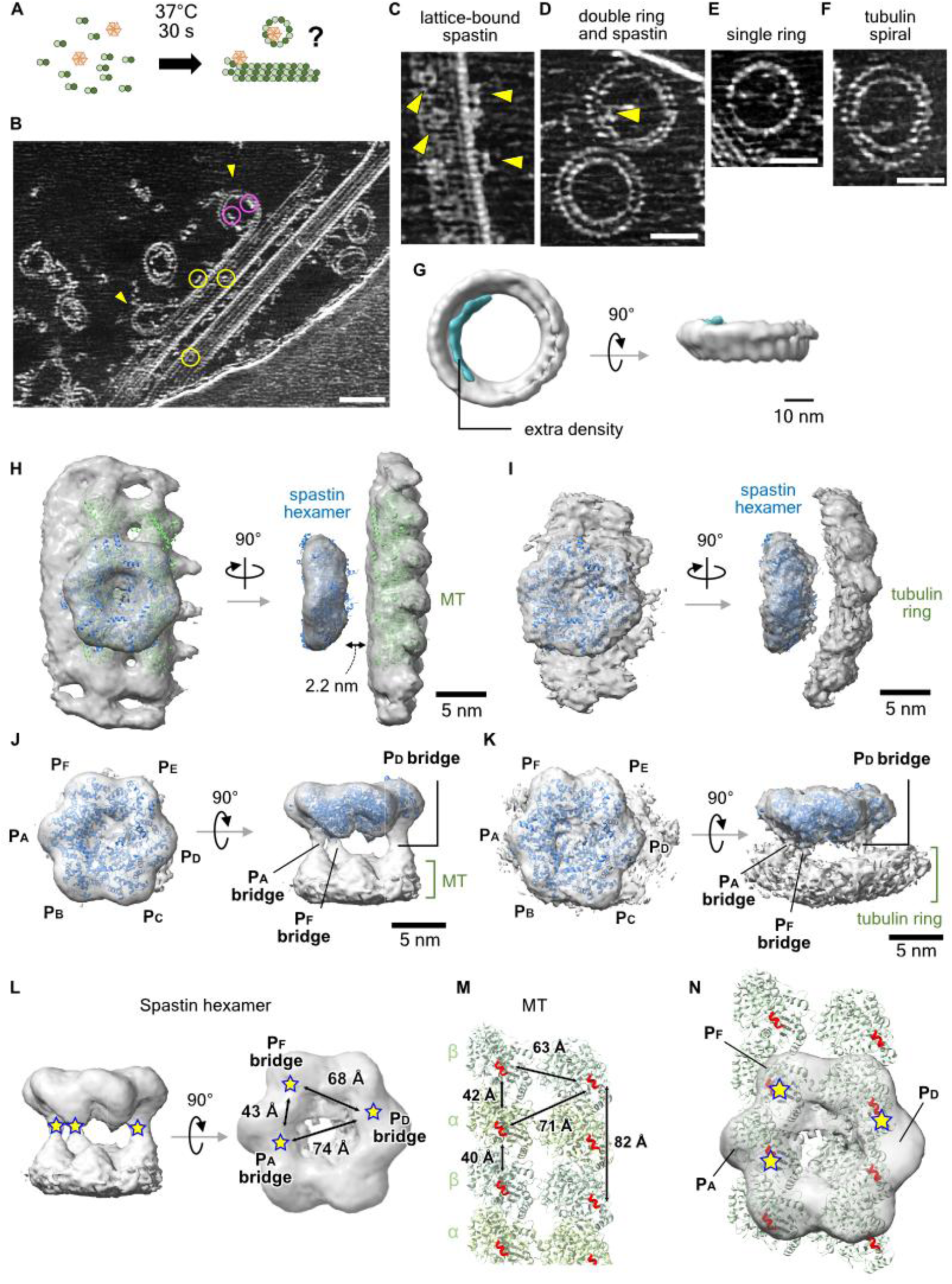
Observation and subtomogram averaging of spastin-tubulin interactions. (**A**) Sample preparation for the cryo-ET of the spastin-dependent microtubule nucleation process. (**B**) A representative denoised, reconstructed and max-projected tomogram. Yellow circles, lattice-bound spastin; magenta circles, ring-bound spastin; arrow heads, tubulin ring incorporation events. Scale bar, 50 nm. (**C–F**) Representative projection images of complexes formed by tubulin and/or spastin. Arrowheads point at spastin hexamers. Scale bars, 25 nm. (**G**) A subtomogram average of the tubulin ring. (**H**) A subtomogram average of lattice-bound spastin hexamers, viewed from two directions. Atomic models of the human spastin hexamer (blue, PDB: 6PEN) and the microtubule lattice (green, PDB: 6DPV) are fitted to the density. (**I**) A subtomogram average of ring-bound spastin hexamers, viewed from two directions. An atomic model of the human spastin hexamer (blue, PDB: 6PEN) is fitted to the density. (**J**) A refined map of the lattice-bound spastin focused on the spastin hexamer, viewed from two directions. Densities that correspond to the interaction of spastin MTBD and tubulin CTT are visible for spastin protomer P_A_, P_D_ and P_F_. (**K**) A refined map of the ring-bound spastin focused on the spastin hexamer, viewed from two directions. spastin protomer P_A_, P_D_ and P_F_ are labeled as in (J). (**L**) The dimensions of the spastin N-terminal densities. The spastin N-terminal bridge densities of P_A_, P_D_ and P_F_ are marked as stars. (**M**) The dimensions of the tubulin CTTs. The tips of tubulin CTTs are colored red. (**N**) The proposed binding mode of the lattice-bound spastin. The map of (L) is overlaid on the model of (M).

Tubulin ring structures could be classified into three types: self-closed single rings, stacked double rings, and spirals (Fig. 4, D–F). Variations in tubulin winding number likely account for the “thin” and “thick” rings observed by negative-stain EM (Fig. 1, F and G). Occasional ring-incorporation events were also detected (Fig. 4B). Although cryo-ET provides static snapshots without temporal resolution, the short 30 s incubation at 37 °C makes it unlikely that these rings reflect depolymerization intermediates; catastrophe frequency during the first 30 s of polymerization is extremely low (<5% at 6 µM tubulin (*66*)). Thus, the rings observed here are more likely polymerizing intermediates. Similar ring structures and incorporation events were observed upon co-incubation with CLIP-170, consistent with the functional overlap with spastin (*49*).

To obtain higher-resolution images of tubulin rings and the spastin-tubulin interaction, we performed subtomogram averaging using RELION 5 (*67*) (fig. S11). We first manually picked ∼350 tubulin rings from denoised tomograms. The refined map revealed two rings of different diameters stacked together (Fig. 4G). Due to the missing wedge, we could not unambiguously determine whether these layers represent two individual rings or a spiral, both of which were observed in the tomograms (Fig. 4, D and F). Nonetheless, additional density within the ring lumen was consistent with bound spastin hexamers, implicating spastin directly in ring assembly.

To further examine the mechanism of ring formation, we next performed subtomogram averaging of the spastin-tubulin complex. We first focused on microtubule lattice-bound spastin to define its binding mode. Approximately 5,500 particles were manually picked to generate an initial model. After iterative refinement, the reconstruction clearly resolved a spastin hexamer bound to the microtubule lattice, with the central AAA domain pore positioned between two protofilaments (Fig. 4H). Although densities corresponding to the spastin MTBD and tubulin CTT were disordered, the spastin hexamer density lay ∼2.2 nm above the microtubule surface, indicating the presence of a spacer between them (Fig. 4H).

We then analyzed ring-bound spastin. From ∼500 manually picked particles, a spastin hexamer was reconstructed on the concave side of the tubulin ring (Fig. 4I). Again, a nanometer-scale gap separated the spastin hexamer from the tubulin ring. Notably, the orientation of the spastin hexamers relative to the tubulin protofilaments was similar in both microtubule–bound and ring–bound reconstructions, suggesting a shared binding geometry. Previous cryo-electron microscopy (cryo-EM) single particle analysis of free spastin in solution showed a spiral spastin hexamer (*13*, *14*). In contrast, binding to either the microtubule lattice or the tubulin ring appears to close the spiral, enforcing a symmetric ring-shaped hexameric conformation.

To gain deeper insight into the configuration, we conducted focused refinement with a mask on the spastin hexamer that partially overlapped the tubulin density. This analysis revealed three asymmetric bridging densities connecting spastin protomers to the microtubule lattice. Based on their positions and our biochemical data (Fig. 3), these contacts are consistent with interactions between the spastin MTBD and the tubulin CTT (Fig. 4J). A similar set of bridging densities was also observed in the ring-bound form of spastin (Fig. 4K; fig. S12A). We designated the six spastin protomers as P_A_ – P_F_ following established convention (*13*), noting that P_A_ was arbitrarily assigned because the protomers are indistinguishable in our hexameric ring conformation.

To understand how these bridges were formed, we measured the dimensions of the bridge densities in the lattice-bound form (Fig. 4J), as the ring-bound densities were too noisy to determine the bridge centers reliably. Considering the relative orientation of spastin hexamer and the microtubule lattice (Fig. 4H), the most plausible interaction pattern is that the P_A_–P_F_ pair bridges longitudinally adjacent tubulin subunits, while the P_D_–P_F_ pair spans neighboring protofilaments (Fig. 4, L and M; fig. S12B). To exclude alternative binding modes, we performed an exhaustive computational search across possible translations and rotations of the spastin hexamer (fig. S13). The best-scoring pose was consistent with our proposed interaction pattern.

In this conformation, the spastin hexamer reinforces lateral contacts between neighboring α- or β-tubulin subunits. Because lateral interactions between protofilaments are a key determinant of nucleation efficiency–for example, tubulin acetylation weakens lateral contacts and thereby inhibits nucleation (*48*)–the P_D_P_F_ bridge is expected to promote efficient microtubule nucleation (Fig. 4N). Consistent with our biochemical assays, which demonstrated that β-CTT is dispensable for nucleation activity, the P_D_P_F_ bridge may engage neighboring α-tubulin subunits (Fig. 3, G–I).

Notably, the central pore of the spastin hexamer lies between protofilaments and is spatially distant from any tubulin CTTs, suggesting that tubulin-binding during nucleation does not require the pore-CTT interaction. Additional conformational changes would be necessary for spastin to sever microtubules if a tubulin CTT were to pass through the pore (*68*). This conclusion is supported by our biochemical assays showing that a pore mutant (A457E) had no effect on ring formation and nucleation (Fig 3, D and E; fig. S8C).

It remains possible that protomers not involved in this trivalent interaction (P_B_, P_C_, and P_E_) contribute to microtubule bundling (figs. S6, A and B, and S10). However, we did not observe any spastin-mediated bridges between microtubules in the reconstructed tomograms, and further investigation will be required to address this possibility. Taken together, these results demonstrate that the spastin hexamer forms trivalent interactions on the microtubule lattice that simultaneously reinforce longitudinal and lateral tubulin contacts, and that a similar mode of interaction underlies assembly of tubulin ring structures.

### Cryo-EM structure of the spastin-induced tubulin ring stack

Our cryo-ET analysis revealed that spastin hexamers bind to the inner side of the rings. However, the orientation of individual tubulins and the mechanism by which tubulin rings of different diameters interact stably despite their symmetry mismatch remained unclear (Fig. 4G). To address these questions, we conducted single particle analysis (SPA) using cryo-EM. To prepare a homogeneous population of rings, we used the non-hydrolyzable GTP analog GTPγS to suppress microtubule formation, thereby arresting the tubulin polymerization at the ring formation step even at 30°C. This treatment increased the population of thick rings, as confirmed by negative staining (Fig. 5A). During cryo-EM data collection, tubulin rings exhibited severe orientation bias, similar to that seen in negative-stain micrographs (Figs. 1, F and G, and 5A). We overcame this problem by adding the non-ionic surfactant 3-[(3-Cholamidopropyl)dimethylammonio]-2-hydroxypropanesulfonate (CHAPSO) (*69*).

**Fig. 5.**
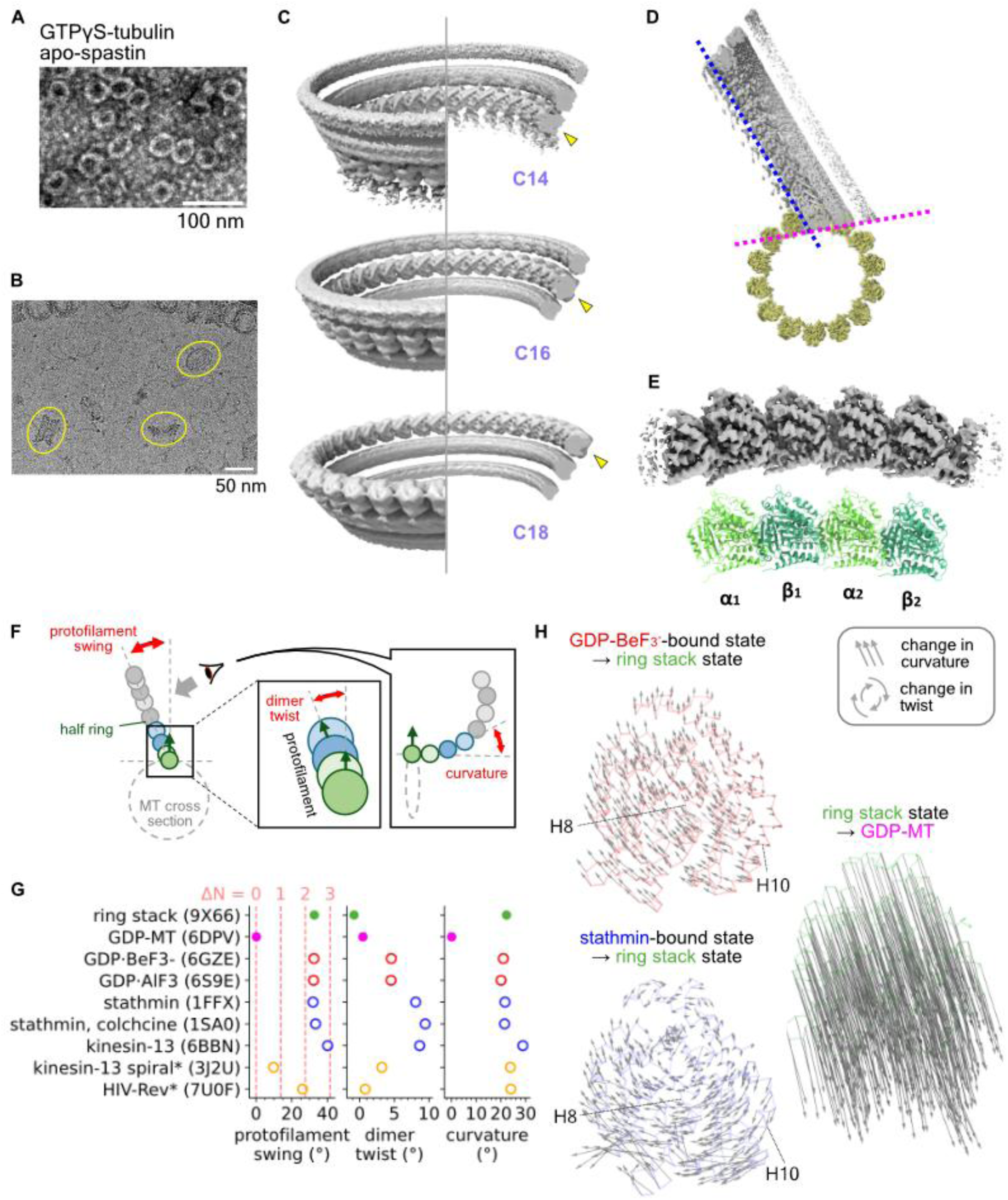
Cryo-EM structure of the spastin-tubulin ring stack. (**A**) Negative-stain micrograph showing that similar tubulin rings are formed with GTPγS instead of GTP. (**B**) A representative cryo-EM micrograph with truncated-cone shaped tubulin ring stacks. (**C**) C14, C16 and C18 reconstruction of the tubulin ring stacks. Each arrowhead indicates the tubulin ring focused for refinement with circular symmetry. (**D**) Tubulin ring stack fitted to the atomic model of microtubule, viewed from plus-to-minus-end direction. Microtubule density map (EMD-7974) is also fitted for the reference. Blue line shows the protofilament swing of the C14 ring. Magenta line corresponds to the plane formed by the lateral contact between protofilaments. (**E**) The 3.41- Å map and the atomic model of the tubulin tetramer in the C16 ring. α_1_, β_1_, α_2_ and β_2_ are labeled from the minus end to the plus end. (**F**) Schematic illustration of the parameters that describe the structure of tubulin tetramers. The arrow pointing from tubulin (circle) is the vector perpendicular to the microtubule surface. (**G**) Scatter plot of the structural parameters calculated from the atomic models of tubulin tetramers as shown in (F). Markers are colored by the type of tetramers. Red: polymerizing state, blue: depolymerizing state, magenta: microtubule, green: tubulin ring stack, and orange: others. Asterisk in the y-axis tick labels means that the atomic models are fitted to the corresponding density map because the deposited PDB file only contains one set of αβ-tubulin (see Method details). Numerical values and the source PDB IDs are listed in Table S2. For the plot of protofilament swing, each dotted line indicates the theoretically allowed values for the corresponding dimer increment numbers (Δ*N*) calculated from the tubulin ring stack formed by WT spastin, detailed in Figs. S18F–S18H. (**H**) Displacement vector maps of β_2_ tubulin, viewed parallel to the protofilament axis. Only the Cα chain of the β_2_ tubulin where vectors start is overlaid. Before displacement calculation, two tetramer models were fitted by α_1_ and β_1_ chain. In this view direction, change in the curvature appears as the linear component, while change in the twist appears as the rotational component, as indicated in (F).

Consistent with cryo-ET reconstructions (Fig. 4, B, D and G), tubulin rings assembled into truncated-cone-shaped stacks, with ring diameters increasing progressively along the stack (Fig. 5B). Under our conditions, most stacks comprised three rings, and pairs of stacks frequently associated via their smallest rings (Fig. 5B). Symmetry-free initial reconstruction indicated 14, 16, and 18 tubulin dimers per ring. Focused refinement with the corresponding symmetry yielded well-defined tubulin densities at 4.16 Å, 4.64 Å, and 7.12 Å, validating these assignments (Fig. 5C; fig. S14).

Tubulin polarity was parallel between rings (fig. S15, A and B), except at the C14-C14 interface of double stacks (fig. S15C). Despite symmetry mismatch and associated lateral registry variations––features likely to generate heterogeneous spastin hexamer binding––we reconstructed a hexameric spastin ring density within the C16 rings, consistent with inner-surface binding observed by cryo-ET (fig. S15D).

To assess the robustness of the ring-stack architecture, we reconstituted tubulin ring stacks using the ATP-state E442Q mutant (fig. S16). Double ring stacks were absent, but high-resolution reconstructions were obtained for C13, C15, and C17 rings (6.07 Å, 4.27 Å, and 5.20 Å, respectively; fig. S16, B and C). Since both apo-wildtype and ATP-E442Q spastin nucleated microtubules (Fig. 3B), we conclude that ring stacks increasing in two-dimer increments are a common architectural motif of the microtubule nucleus, whereas the exact ring size is not fixed.

To relate the SPA structures to microtubule geometry, we fitted the C14 ring into a published microtubule model (PDB: 6DPV (*70*)) (Fig. 5D). The protofilament trajectory showed a counterclockwise tilt (“swing”), in agreement with counterclockwise tilts at polymerizing and depolymerizing ends observed by cryo-ET and predicted by molecular-dynamics simulations (*44*, *71*, *72*). The inter-ring interface lay approximately tangent to the microtubule surface (Fig. 5D), providing a natural geometric explanation for the truncated-cone architecture, in which progressively larger rings preserve tangential contacts during stacking (fig. S15E).

### Intrinsic swing angle of tubulin favors ring stacking

Although the spastin density was blurred due to symmetry mismatch and flexibility, our reconstructions captured a tubulin complex consistent with an intermediate in spontaneous nucleation. To probe the structural basis of this intermediate, we performed focused refinement on the tubulin tetramer within the C16 ring, resulting in a 3.41-Å global resolution map (Fig. 5E; fig. S17, A and B; Table S1). At this resolution, secondary structures, α- and β-tubulin identity, and nucleotides were clearly resolved (fig. S17, C–E).

Periodic tubulin assemblies impose strong conformational constraints (*73*), yet the protofilament swing in our ring stack (∼30°) closely matched that of non-polymerized tubulin tetramers, detailed below (Fig. 5, F and G; fig. S18, A–E). Using intrinsic spacings—the longitudinal inter-dimer distance 82 Å (*74*) (Fig. 4M) and the lateral interval measured from the C1 map (fig. S18, F and G) —we calculated the swing angles allowed for different dimer increments (Δ*N*). With the measured lateral spacing and Δ*N* = 2 (applicable to WT and E442Q), the predicted swings were 27.6° and 28.1° (fig. S18H), closely matching the intrinsic tetramer value from atomic models (Fig. 5G). Alternative solutions (Δ*N* = 1: 13.8° for WT and 14.0° for E442Q, Δ*N* = 3: 41.4° for WT and 42.1° for E442Q) deviated substantially. These results indicate that tubulin possesses an intrinsic swing angle that is geometrically optimal for ring stacking, which explains why the swing angle remains essentially constant across ring, polymerizing, and depolymerizing states.

### Ring stacks promote microtubule formation by inter-dimer untwisting

To assess how the tetramer is configured within the ring stack, we compared it with tetramers from polymerizing ends (tubulins bound to GTP analogs in the exchangeable site: 6GZE, 6S9E), depolymerizing ends (tubulins complexed with microtubule depolymerizers: 6BBN, 1FFX, 1SA0), and the microtubule lattice (6DPV), using geometric descriptors including protofilament swing, dimer twist, and curvature (Fig. 5F; fig. S18, A–E). Protofilament swing angles clustered near 30° for both tubulin tetramers and our ring structure, except for the kinesin-13-bound tetramer that has exceptionally large protofilament swing (Fig. 5G; Table S2). Similarly, overall curvature remained unchanged during ring formation (Fig. 5G), consistent with the match between the curvature of rings and that of protofilament tips at polymerizing and depolymerizing ends (*44*) (Fig. 1, F and G).

By contrast, dimer twist clearly distinguished the structural states: it was largest (∼9°) at depolymerizing ends, ∼4.5° at polymerizing ends, but nearly 0° in both the ring stack and the microtubule lattice (Fig. 5G). This indicates that the ring formation enforces a microtubule-type, non-twisted interface during the transition from soluble tubulin to the lattice. Because dimer twisting displaces protofilaments azimuthally, the large dimer twist observed at depolymerizing ends likely drives spiral formation (*43*, *75*). Consistent with this interpretation, the tubulin ring induced by HIV Rev (*76*) is essentially untwisted, whereas tubulin spirals formed outside the microtubule-kinesin-13 complex (*77*) show weak twisting (Fig. 5G).

To confirm that the difference in the dimer twist parameter reflects true rotational motion, we fitted tetramer models by aligning the first tubulin dimer and calculated the Cα displacement vector map for the second β-tubulin (*78*). When viewed parallel to the protofilament axis, the conformational change from the polymerizing/depolymerizing states to the ring stack state contained a clear clockwise rotational component, whereas the conformational change from the ring stack state to the microtubule lattice is essentially linear (Fig. 5H). A similar rotational component was evident in the displacement vector maps from polymerizing/depolymerizing states to the microtubule lattice (fig. S19A).

To pinpoint the structural basis of the twist differences, we performed RMSD analysis among the tetramers. The deviations between polymerizing/depolymerizing states and the ring stack state were concentrated at the inter-dimer interface, particularly in helices H8 and H10 of α_2_-tubulin (fig. S19B). These results suggest that spastin resolves the inter-dimer twisting by reorganizing these regions, thereby switching tubulin from a depolymerization-prone trajectory toward a nucleation-competent state. Collectively, these data define inter-dimer untwisting as the hallmark conformational change that underlies tubulin ring stack formation and microtubule nucleation.

## Discussion

Our work identifies human spastin as an ATPase-independent microtubule nucleator and bundler. Biochemical assays, negative staining EM, and fluorescence microscopy imaging demonstrate that, in the presence of free tubulin, spastin increases microtubule mass primarily through spontaneous nucleation rather than severing. Subtomogram averaging and single particle analysis further provide a structural rationale for this activity: spastin hexamers bridge lateral and longitudinal tubulin contacts to assemble ring stacks, thereby stabilizing nucleation-competent intermediates that promote microtubule formation.

Across TIRFM assays––monitoring severing (Fig. 1, C and D), nucleation (Fig. 2, C–E), and microtubule dynamics (Fig. 2, F and G) –– soluble tubulin strongly suppressed severing, whereas spastin-driven nucleation and bundling remained robust over a broad concentration range. These observations can be explained by tubulin ring formation mediated by the interactions between spastin MTBD and the tubulin CTT. Tubulin ring formation decreases the pool of spastin molecules available for severing and ultimately reduces the severing rate below the lattice repair rate (fig. S2). We therefore propose that, under physiological conditions, spastin functions primarily as a microtubule network architect rather than a breaker. Marked severing in cells may occur only under conditions of tubulin depletion and/or elevated spastin levels, reconciling the severing-dominant behavior in tubulin-free reconstitutions with the assembly-promoting roles observed here (Fig. 6A).

**Fig. 6.**
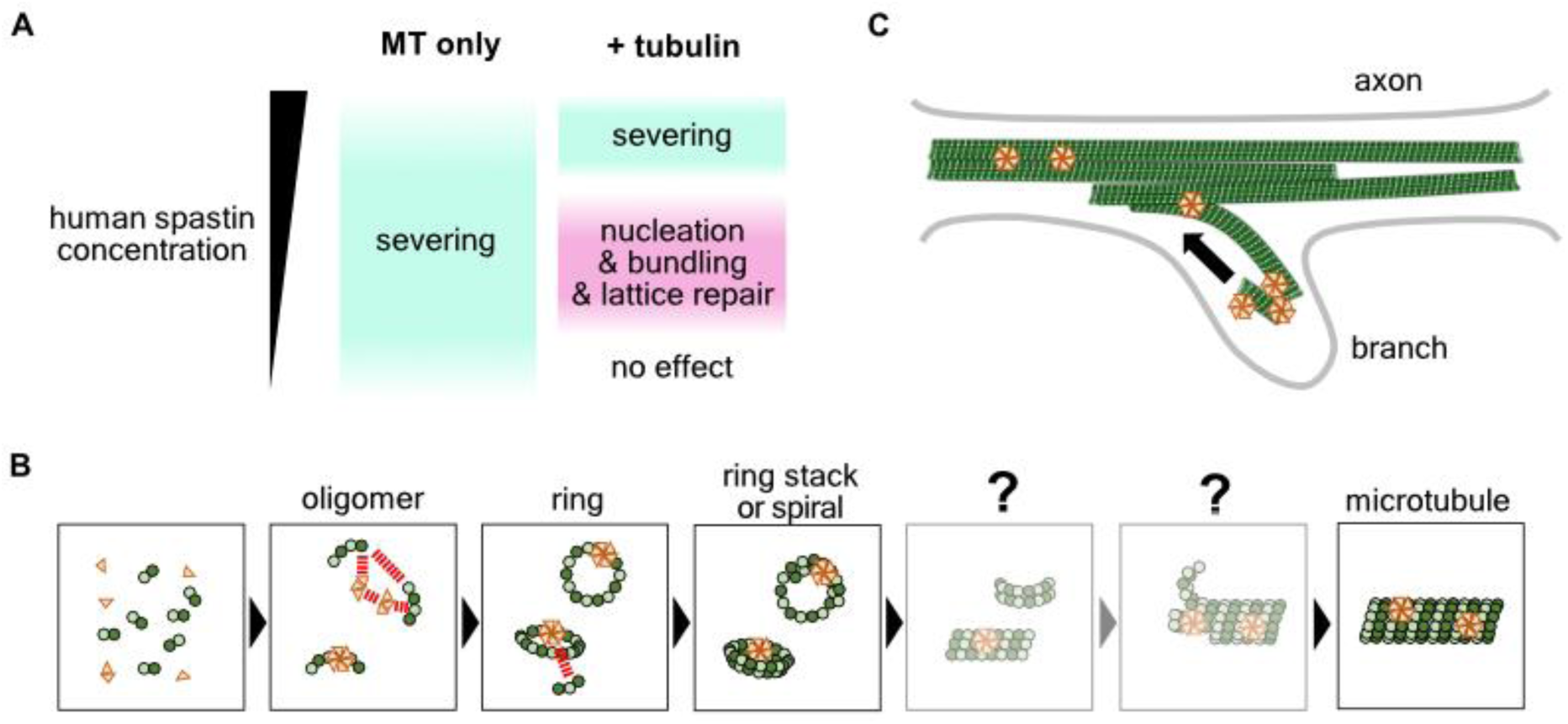
Model of microtubule nucleation and reorganization by human spastin. (**A**) The scale of the spastin activities. Free tubulin has inhibitory effect on the spastin severing activity because it forms spastin-tubulin rings and repairs lattice damages induced by spastin. As a result, the severing activity stands out in the reconstitution assays that only use polymerized microtubules, but much higher concentration of spastin is needed for the severing activity to outcompete the nucleation activity, bundling activity and lattice repair events in the presence of free tubulin. (**B**) Schematic illustration of the model of the microtubule spontaneous nucleation with spastin. Iterative tubulin oligomerization in the longitudinal direction forms individual tubulin rings. Oligomerization of spastin and oligomerization of tubulin may allosterically facilitate each other, resulting in a switch-like response of the ring-formation process. The lateral contacts are formed by either stacking rings with different diameter, or spiral formation, which can also be promoted by the spastin hexamer bound inside the ring. Transition from ring stack or spiral to microtubule requires the breakage of the tubulin ring and formation of microtubule template, but these steps could not be directly observed in this study. Because spastin uses the same binding mode for both tubulin ring and microtubule lattice, dissociation of spastin is not needed for microtubule formation. (**C**) Schematic illustration of the microtubule network reorganization in axons by spastin. Microtubules nucleated by spastin can directly be connected to the existing microtubule arrays in the axon by bundling, by which axon branches will be stably formed.

Importantly, our findings also address a long-standing question in cytoskeletal biology: whether tubulin rings represent intermediates on the path to depolymerization or polymerization. Historically associated with catastrophes and protofilament peeling, rings have been interpreted primarily as disassembly products (*45*, *46*). Our data instead reveal that rings can represent two structurally and functionally distinct states. In the absence of regulatory factors, tubulin oligomers tend to form twisted spirals that bias assemblies toward depolymerization. In contrast, spastin converts these intrinsically twisted intermediates into straightened, lattice-like ring stacks that are competent for microtubule growth. This duality in ring fate is also supported in vivo: recent in situ cryo-ET imaging of regenerating axon shows that tubulin rings accumulate at polymerizing sites and participate in rapid microtubule polymerization (*79*). Together, these observations indicate that tubulin rings are not obligate depolymerization intermediates but bifurcation points whose fate is determined by regulators such as spastin.

The structural and geometric features of the intermediates help explain why spastin is required to redirect rings toward polymerization (Fig. 6B). Prior studies have placed tubulin oligomers at the earliest stage of spontaneous nucleation (*48*, *60*). We find that such oligomers can close into rings; however, because the longitudinal interface exhibits a non-zero dimer twist (Fig. 5G), closure into a stable ring is likely energetically unfavorable. This is consistent with the requirement for significantly higher tubulin concentrations for spontaneous nucleation than for elongation (*54*).

Once spastin-induced rings are formed, stacking becomes favorable because the protofilament swing angle is already near the optimal value for ring-ring stacking (Fig. 5G; fig. S18, F and G). We propose that the spastin-tubulin contacts simultaneously strengthen both longitudinal and lateral interactions, thereby driving the otherwise rare oligomer-to-ring transition and stabilizing ring-stack intermediates. Although we did not dissect the subsequent ring-breakage step required to generate microtubules (*80*), binding of spastin should be compatible with this conversion, as spastin does not need to dissociate from the tubulin complex considering the similarity between lattice-bound and ring-bound states (Fig. 4, H–K). Ring-incorporation events observed by cryo-ET (Fig. 4B) may capture features of this conversion.

Spastin also reorganizes filament distribution in ways that could influence the patterning of cellular arrays (Fig. 6C). We observed rapid re-bundling of fragments, pulling-mediated reorientation, and frequent zippering and connection events of elongating filaments (Fig. 2, F and G). These behaviors favor the formation of ordered arrays over randomly scattered fragments and may relate to axon branching and presynaptic polymerization in neurons (*21*, *81*). Notably, tau shares nucleation and bundling activities and is enriched in axons (*58*, *82*–*84*), but tau and spastin interact with microtubules in a mutually exclusive manner (*37*, *85*, *86*). Another microtubule remodeler, SSNA1, co-localizes with spastin at branching points and participates in axonal development, while simultaneously inhibiting spastin-mediated severing (*87*, *88*). Future live-cell studies will be important to clarify how these factors partition remodeling activities in space and time.

Regulation by post-translational modifications (PTMs) likely tunes the balance among severing, nucleation, and bundling. Spastin phosphorylation and SUMOylation influence neuronal morphology (*89–91*), while tubulin polyglutamylation modulates both severing efficiency and neuronal phenotype (*27*, *35*, *92*). Because we used purified spastin and brain-derived tubulin enriched in PTMs (*93*), reconstitutions with defined PTM patterns will be essential to quantify how PTMs redistribute spastin activity across its multiple functional outputs.

Species comparisons further underscore that the strategy of microtubule amplification by spastin is not universal. In Drosophila, DmSpastin primarily amplifies microtubule mass by severing pre-existing polymers to create seeds—a microtubule-dependent amplification route (*31*, *34*, *68*). By contrast, human spastin nucleates microtubules de novo through MTBD-dependent ring stack formation, even without ATP hydrolysis. Drosophila may therefore require tighter regulation of severing activity, or may compensate for weaker microtubule-independent nucleation through an intrinsically higher tubulin nucleation propensity (*94*). These differences caution against direct extrapolation of DmSpastin mechanisms to human HSP.

From a clinical perspective, our findings may inform new therapeutic strategies for HSP. The prevailing loss-of-function model, where HSP symptoms arise from insufficient microtubule severing, has been widely used to explain the effects of individual spastin mutations. However, this framework does not fully account for the clinical spectrum of HSP, including the relatively mild impact of certain missense mutations in humans and reported gain-of-function phenotypes (*26*). Our results suggest that spastin variants may differentially nucleate and organize microtubules, highlighting the need to assess pathology in a more multifaceted and comprehensive manner. These insights ultimately open the possibility for more effective and personalized therapeutic approaches for HSP.

## Acknowledgments

During the preparation of this work, the authors used ChatGPT and DeepL to improve phrasings. We are grateful to Dr. Shinsuke Niwa (Tohoku University) for kindly providing the human *spastin* gene. We thank K Chin for the research management support, and the member of Nitta lab for the discussion. The cryo-EM experiments were performed at SPring-8 with the approval of the Japan Synchrotron Radiation Research Institute (JASRI) (Proposal No. 2021B2536, 2022A2536, 2022B2542, 2023A2542, 2023B2537, 2024B2531, 2024B2532) and AMED-BINDS JP25ama121001.

## Funding

Japan Society for the Promotion of Science KAKENHI grant JP21H05254 (RN)

Japan Society for the Promotion of Science KAKENHI grant 22K06809 (TI)

Japan Society for the Promotion of Science KAKENHI grant 25H02379 (TI)

Japan Society for the Promotion of Science KAKENHI grant 23KJ0472 (HL)

JST Moonshot Research and Development Program grant JPMJMS2024-7 (RN)

JST FOREST JPMJFR214K (TI)

Japan Agency for Medical Research and Development (AMED) CREST grant JP21gm161003 (TI)

The Hyogo Science and Technology Association (RN)

## Author contributions

Conceptualization: HL, TI, RN

Methodology: HL, NM, ToK

Investigation: HL, NM, ToK, TaK, HS, SK, EN

Visualization: HL

Funding acquisition: TI, RN

Project administration: RN

Supervision: TI, RN

Writing – original draft: HL

Writing – review & editing: HL, TaK, HS, TI, RN

## Data and materials availability

Plasmids generated in this study will be shared upon request. Raw data of TIRFM, negative-stain EM, turbidity assay and electrophoresis, reconstructed cryo-EM maps, and codes for statistical testing, analysis and plotting are deposited on Zenodo with DOI 10.5281/zenodo.17084043. Electron density maps and the atomic models are deposited on Electron Microscopy Data Bank and Protein Data Bank under accession number as follows. Subtomogram averages: ring stack (EMD-66612), lattice-bound spastin (EMD-66613, EMD-66614), and ring-bound spastin (EMD-66615, EMD-66616). Single particle analysis: tubulin tetramer in the ring stack (EMD-66608, PDB: 9X66), tubulin ring stack reconstructed with C14, C16 and C18 symmetry (EMD-66609, EMD-66610, EMD-66611).

## Supplementary Materials

### Materials and Methods

#### Tubulin Purification and Labeling

Tubulin was purified from porcine brains by two cycles of polymerization and depolymerization (Castoldi and Popov, 2003) and dissolved in PEM (100 mM PIPES-KOH, 1 mM EGTA, 2 mM MgSO_4_, pH 6.9) supplemented with 0.1 mM GTP. The tubulin concentration was measured by the 280-nm absorbance using the extinction coefficient 0.115 µM^-1^cm^-1^ (*1*). For preparation of GTPγS tubulin, tubulin was polymerized with 1 mM GTP, and the microtubule pellet was resuspended and depolymerized in PEM containing 5 mM GTPγS (NU-412-20, Jena Bioscience). For preparation of GMPCPP tubulin, tubulin was polymerized with 0.1 mM GMPCPP (NU-405L, Jena Bioscience), and the microtubule pellet was resuspended and depolymerized in PEM containing 0.1 mM GMPCPP. For labeling, polymerized microtubules were mixed with AF 568 NHS ester (#14820, Lumiprobe), AF 647 NHS ester (#16820, Lumiprobe) or Biotin-LC-LC-NHS (#B6096, Tokyo Chemical Industry) dissolved in dimethyl sulfoxide (DMSO), and non-reacted dye was removed by two cycles of centrifugation/resuspension or by the polymerization/depolymerization cycle during the abovementioned tubulin purification.

#### Protein Purification

Glutathione *S*-transferase (GST)-tag-conjugated human spastin (a.a. 228-) and its mutants were expressed and purified from Rosetta(DE3) as follows. Cells were cultured in LB media at 37°C until OD_600_ reaches ∼0.6 and chilled on ice to prevent further proliferation. Isopropyl β-D-thiogalactopyranoside (IPTG) was added to the culture to the final concentration 0.1 – 0.2 mM for the induction of protein expression, and the cells were further cultured at 25°C overnight. Cells were collected by centrifugation, frozen with liquid nitrogen, and stored at −80°C until use. For protein purification, cells were resuspended in lysis buffer (50 mM HEPES, 1 M NaCl, 2.5 mM MgCl_2_, pH 7.5 adjusted with KOH) supplemented with 5 mM 2-mercaptoethanol and protease inhibitor cocktail (0.66 μM Leupeptin, 2 μM pepstatin A, 2 mM benzamidine, 1 mM phenylmethylsulfonyl fluoride), and lysed by sonication. Cell debris were cleaned by centrifugation at 16,000 g, 4°C for 10 min. The supernatant was collected and incubated with Glutathione Sepharose™ 4B resin (17075605; cytiva) equilibrated with lysis buffer for 1 h. Resin was washed with more than 3× volume of lysis buffer and the GST-tag was cleaved by 3C protease. The cleaved fraction was concentrated by Amicon Ultra-4 centrifugal filter (UFC801096; Millipore) and further purified by size-exclusion chromatography using Superose™ 6 Increase 10/300 GL (29091596; cytiva) with sizing buffer (20 mM HEPES, 150 mM NaCl, 2.5 mM MgCl_2_, 2 mM DTT, pH 7.5 adjusted with KOH) at flow rate 0.8 mL/min. Peak fractions were concentrated by Amicon Ultra-4 centrifugal filter. Protein concentration was measured by Micro BCA Protein Assay Kit (23235; ThermoFisher Scientific).

6×His-tag-conjugated mouse rigor KIF5C mutant (a.a. 1 – 411, G235A) was expressed and purified from Rosetta(DE3) as follows. Cells were cultured in LB media at 37°C until OD_600_ reaches ∼0.6 and chilled on ice to prevent further proliferation. IPTG was added to the culture to the final concentration 0.5 mM and the cells were further cultured at 25°C for 4 h. Cells were collected by centrifugation, frozen with liquid nitrogen, and stored at −80°C until use. For protein purification, cells were resuspended in lysis buffer (50 mM Tris, 100 mM NaCl, pH 8.0 adjusted with HCl) supplemented with protease inhibitor cocktail, and lysed by sonication. Cell debris were cleaned by centrifugation at 16,000 g, 4°C for 15 min. The supernatant was collected and incubated with HIS-Select Nickel Affinity Gel (P6611; Millipore) equilibrated with kinesin lysis buffer for 30 min. Resin was washed with 3× column volumes of lysis buffer supplemented with 10 mM imidazole, and further washed with 3× column volumes of lysis buffer supplemented with 20 mM imidazole. Protein was eluted with lysis buffer supplemented with 300 mM imidazole and concentrated with Amicon Ultra – 0.5 mL centrifugal filter (UFC501096; Millipore). Protein concentration was measured by absorbance at 280 nm.

#### Microtubule Turbidity Assay

Nucleation and polymerization of tubulins were monitored by absorbance at 350 nm using spectrophotometer (DS-11+, DeNovix) as follows. Mixtures containing 15 µM tubulin were prepared in PEM buffer supplemented with 1 mM GTP. Spastin, 10 µM taxol and 1 mM ATP were also added depending on the condition. Tubulin and spastin was added to the mixtures in the final step so that tubulin-spastin complex will not unexpectedly be formed due to the temporary high concentration during preparation. The mixtures were immediately put in a cuvette and inserted into the spectrophotometer prewarmed to 37°C. Absorbance was measured every 30 second for 30 min.

#### Limited Proteolysis by Subtilisin

For preparation of β-CTT-digested tubulin samples (S_β_-tubulin), 100 μM tubulin in PEM and 1 mM GTP was first polymerized by incubation at 37°C for 30 min and stabilized by adding half volume of glycerol. Subtilisin (P5380, Sigma) was added to this microtubule solution by mixing 2.5 mg/mL subtilisin dissolved in PEM 50% glycerol and incubated at 37°C for 45 min. The weight ratio of subtilisin to tubulin is adjusted to 2%. Proteolysis was terminated by adding final 40 mM of phenylmethylsulfonyl fluoride (PMSF) dissolved in isopropanol. Polymerized microtubules were purified by ultracentrifugation at 49,000 rpm, 35°C for 12 min using TLA-100.3 rotor (349481; Beckman). Microtubule pellet was resuspended in iced PEM and depolymerized by pipetting. Aggregation and non-depolymerized microtubules were removed by centrifugation at 20,000 g, 4°C for 10 min. Proteolysis was confirmed by the band shift in SDS-PAGE.

#### Tubulin Ring Pelleting Assay

To form tubulin rings, 8 μM tubulin and 1.6 μM spastin were co-incubated in 1.5-mL ultracentrifuge tubes on ice for 20 min in the total volume of 25 μL unless otherwise stated. For the negative control, tubulin was incubated without spastin. Mixtures were subsequently ultracentrifuged at 54,800 rpm (200,000 g), 4°C for 20 min using S55A2 rotor (himac). After aliquoting 6 μL of the supernatant for SDS-PAGE, all the supernatant were removed and the pellets were rinsed with 25 μL of BRB80 (80 mM PIPES, 1 mM EGTA, 2 mM MgSO_4_, pH 7.3) three times. The pellets were then resuspended in BRB80 12.5 μL, and 6 μL was aliquoted for SDS-PAGE. SDS-PAGE was conducted with 9% polyacrylamide gels. After electrophoresis, gels were stained with CBB for 20 min and destained by water. Band and background intensities were measured by the rectangle tool of ImageJ (NIH). Binding ratio was calculated as (*I_P_* − *I_BG_*)/(2(*I*_*S*_ − *I_BG_*) + *I_P_* − *I_BG_*), where *I_P_*, *I*_*S*_ and *I_BG_* are the mean intensity of pellet band, supernatant band, and background, respectively. Supernatant band intensity is multiplied by 2 because 25 µL mixture was resuspended in 12.5 µL after pelleting.

#### Total Internal Reflection Fluorescence Microscopy

Images were acquired using Eclipse Ti2 TIRF microscope (Nikon) equipped with an iXon Life 897 EMCCD camera (Andor), CFI Apochromat TIRF lens (NA 1.49, 100×; Nikon), and lens/stage heaters (Tokai Hit). To make glass flow chambers, 22 mm × 22 mm coverslips (474030-9020-000; ZEISS) were treated in 1 M HCl overnight and sonicated for 30 min three times in deionized water, once in 50% ethanol, 70% ethanol and 95% ethanol. A full-size coverslip and a coverslip cut into 1/4 size with glass cutter was sticked together with double-sided tapes (Nichiban).

##### Severing assay

60 µM 5%-biotinylated-3%-AF568-labeled tubulin in BRB80 containing 1 mM GTP was incubated at 37°C for 10 min, stabilized with 100 µM taxol, and further incubated for 20 min to complete polymerization. Microtubules were pelleted by centrifugation at 20,000 g, 25°C for 25 min and resuspended in BRB80 supplemented with 100 µM taxol. Glass chamber was first treated with PLL-g-PEG-biotin for 5 min and subsequently incubated with 1 mg/mL biotinylated BSA for 2 min. After washing out unbound fraction, 0.5 mg/mL NeutrAvidin was added and incubated for 2 min, and was blocked with 5 mg/mL casein for 2 min. Microtubule was diluted in BRB80 supplemented with 10 µM taxol and added to the chamber. To quantify severing inhibition by free tubulin, non-bound microtubules were washed out with 1x BRB80 10% glycerol, and spastin in glycerol-based severing buffer (BRB80, 10% glycerol, 1 mM ATP, 5 mg/mL casein, 2.5 mM PCA, 1.25 mM Trolox, 0.0015 U/µL PCD) was applied. Images were acquired for 5 min at 2.5-s interval for three positions with stage and lens heater set to 28°C. To confirm severing activity of spastin mutant, microtubules were washed out with BRB80 supplemented with 10 µM taxol, and spastin in taxol-based severing buffer (BRB80, 10 µM taxol, 1 mM ATP, 5 mg/mL casein, 2.5 mM PCA, 1.25 mM Trolox, 0.0015 U/µL PCD) was applied. Images were acquired at 1-s interval for 1 min with 0.2 µM spastin, and subsequently 2 µM spastin unless severing events are already observed. Stage and lens heater were set to 28°C.

The progression of microtubule severing was analyzed by the total fluorescence intensity of individual microtubules. Acquired image stacks were processed by rolling-ball algorithm of 20-pixel radius, drift-corrected and ROIs were manually drawn using the segmented line tool of ImageJ. For Figure S6, polygon ROIs were used instead because the drift cannot be precisely corrected. Measured mean intensity traces were saved as CSV files. For each trace, measured fluorescence intensities were normalized by dividing by the intensity of the first frame.

##### Repairing assay

Taxol stabilized, 5%-biotinylated-3%-AF568-labeled microtubule was immobilized to the surface of glass chamber in the same way as in the severing assay. 8 µM 14%-AF647-labeled tubulin in repairing buffer (BRB80, 10% glycerol, 1 mM ATP, 1 mM GTP, 5 mg/mL casein, 2.5 mM PCA, 1.25 mM Trolox, 0.0015 U/µL PCD) with or without 150 nM spastin was added to the chamber. Images were acquired for 5 min at 10-s interval for three positions with stage and lens heater set to 32°C.

The progression of microtubule severing was analyzed by the total fluorescence intensity of AF647 channel along each microtubule. Acquired image stacks were processed by rolling-ball algorithm of 30-pixel radius, drift-corrected and ROIs were manually drawn using the segmented line tool of ImageJ. Measured mean intensity traces were saved as CSV files for plotting.

##### Depolymerization assay

10 µM 10%-biotinylated-5%-AF647-labeled tubulin in PEM buffer containing 0.1 mM GMPCPP was incubated at 27°C for 1 h and 37°C for 1 h to polymerize the microtubule seeds. Seeds were pelleted by centrifugation at 20,000 g, 25°C for 25 min, and resuspended in BRB80 50% glycerol. Microtubule seeds were immobilized to the glass chamber in the same way as the severing assay. Stage and lens heater were set to 32°C, and 15 µM 5%-AF568-labeled tubulin with 0.5 mM GTP was added to let microtubules polymerize from seeds for 5 min. After washing tubulin out with BRB80 25% glycerol, depolymerizing buffer (BRB80, 5 mg/mL casein, 10% glycerol, 0.1% methylcellulose, 2.5 mM PCA, 1.25 mM Trolox, 0.0015 U/µL PCD) with or without 0.5 µM spastin was applied. Images were acquired at 5-s interval for 5 min. Acquired image stacks were processed by rolling-ball algorithm of 30-pixel radius and drift corrected based on the AF647 channel. Kymographs were calculated for non-overlapping microtubules and the shrinkage rate was measured by the slope of the line ROIs in ImageJ. Because shrinkage rate was not constant and pause events were frequently seen for many microtubules, mean shrinkage rate was measured by drawing a single line ROI for each microtubule, from the starting point to the end point of depolymerization.

##### Nucleation assay

12 µM rigor kinesin (KIF5C G235A) was added to the glass chamber and subsequently blocked with 5 mg/mL casein. Z focus was determined by autofluorescence of puncta in the solution. Subsequently, mixture containing 15 µM 5%-AF568-labeled tubulin, 1 mM GTP and 1 mg/mL casein in BRB80 was applied. 1 µM spastin and 1 mM ATP were added depending on the condition. Similar to the turbidity assay, tubulin and spastin was added to the mixtures in the final step. Time-lapse images were acquired in four positions at 10-s interval for 20 min with stage and lens heater set to 32°C. To quantify the increase of microtubule mass, image background was calculated for each frame by median projection along the positional axis and median filter of 20-pixel radius in the xy axes. Mean intensity of each image frame from four positions was calculated for plotting.

##### Microtubule bundling assay

5 µL of 60 µM 5%-AF568-labeled tubulin was polymerized at 37°C for 10 min, stabilized by adding 100 µM taxol, and further incubated at 37°C for 20 min. Microtubules were collected by centrifugation at 20,000 g, 30°C for 25 min and the pellet was resuspended in 10 µL of BRB80 supplemented with 100 µM taxol. This solution was used as 30 µM microtubule stock within a day. Concentration of spastin and taxol-stabilized microtubule was adjusted by BRB80 containing 25% glycerol, mixed together, incubated for 2 min, and directly added to an empty chamber. After 2-min incubation in the chamber, floating proteins were washed out with BRB80 containing 25% glycerol, and images were acquired at 10 positions. Bundling efficiency was quantified by measuring the mean intensity along microtubules as follows. All image slices were first process by Gaussian filter of 1-pixel sigma and rolling-ball algorithm of 30-pixel radius. The threshold value to distinguish microtubule signals from the background was determined to be 500 using the threshold tool of ImageJ. Processed images were binarized and skeletonized to create microtubule mask images. Mean intensity in the mask area was calculated for each image slice.

##### Dynamics assay

Microtubule seeds were immobilized on the glass chamber in the same way as the depolymerization assay. Stage and lens heater were set to 32°C. After washing out non-bound seeds with BRB80, 12 µM 5%-AF568-labeled tubulin with 0, 1 or 2 µM spastin in the dynamics assay buffer (BRB80, 1 mM GTP, 1 mM ATP, 5 mg/mL casein, 0.1% methylcellulose, 2.5 mM PCA, 1.25 mM Trolox, 0.0015 U/µL PCD) was applied. Images were acquired at 10-s interval at three positions for 30 min. The acquired images were processed by rolling-ball algorithm with 30-pixel radius and drift-corrected based on the channel of microtubule seeds. Angles before microtubule zippering or connection were measured by the ImageJ angle ROIs and exported as CSV files for plotting and statistical testing. Microtubules that are completely parallel to each other were excluded from this analysis, because whether they are bundled cannot be determined.

### Negative-stain Electron Microscopy

For negative staining electron microscopy, carbon-coated 200-mesh Cu grid (EM Japan) was glow-discharged prior to preparation of the specimen. Observation was conducted using JEM-1400 Plus electron microscope (JEOL) at acceleration voltage 120 kV and images were obtained with a JEOL Matataki Flash camera.

For samples after turbidity assays, 4 µL of the mixture in the cuvette was directly taken and applied to the grid. After blotting with a filter paper, the grid was stained with 4 µL of 2% uranyl acetate twice.

For visualization of tubulin rings, 8 µM tubulin or 4 µM S_β_-tubulin were incubated with equimolar of spastin in BRB80. 1 mM AMPPNP, 1 mM ADP or 1 mM ADP + 2 mM Na_3_VO_4_ was added to this mixture depending on the condition. This mixture was incubated at 37°C for 30 min, and 4 µL was added to a grid. After blotting with a filter paper, the grid was stained with 4 µL of 2% uranyl acetate twice.

### Cryo-electron Tomography

A glow-discharged holey carbon grid (R1.2/1.3, Cu, 300mesh; Quantifoil) was first made as hydrophilic its surface by applying 0.01% NP-40 aqueous solution. Mixture of 30 µM tubulin, 30 µM spastin, 0.01% NP-40, 1 mM GTP, and 10-nm gold (CGM5K-10-50, cytodiagnosis) was incubated at 37°C for 30 s, and 4 µL of it was placed on the grid inside the blotting machine (Vitrobot Mark IV; Thermo Fisher) kept at 37°C and 100% humidity. Grids were immediately blotted and plunged in liquid ethane. Data collection was performed on Glacios microscope (Thermo Fisher) equipped with Falcon 4i Direct Electron Detector (Thermo Fisher) using Tomography 5 software (Thermo Fisher). Tilt series were obtained from −51° to 51° with 3° increment, with the total dose of ∼110 e^-^/Å^2^, 200 kV acceleration voltage, and target defocus value ranging −2 to −4 µm. 45,000× magnification was used, which resulted in 3.1 Å/pixel.

Collected movies were analyzed using RELION tomography toolkit (*2*) as outlined in Figure S11. Movies were first motion-corrected with odd/even split, CTF-corrected, and after bad image slices were manually excluded, tilt series were automatically aligned using gold beads as fiducials. Gold beads were detected by difference-of-Gaussian filter and thresholding, and erased by substitution with white Gaussian noise by executing a Python script as a RELION job. The gold-erased tilt series were used for reconstruction of half tomograms with 7 Å/pixel scale, and three of them containing representative protein densities were used for training cryo-CARE denoiser. All the tomograms were denoised, and correctness of the tilt angle handedness is confirmed at this point by the microtubule lattice pattern. Particles of tubulin rings, lattice-bound spastin hexamers and ring-bound spastin hexamers were manually picked using napari (*3*) and pseudo subtomograms were extracted as 2D image stacks. Initial models were built by 4× or 2× binned subtomograms and the final auto-refinement jobs were performed by re-extracted subtomograms without binning. Masks for refinement were manually drawn using the 3D painting tool of napari or the volume eraser tool of ChimeraX (*4*). Atomic models of spastin hexamer (PDB: 6PEN) and undecorated GDP-MT (PDB: 6DPV) were fitted to the maps by the fitmap command of ChimeraX.

### Cryo-electron Microscopy and Single Particle Analysis

Tubulin with 5 mM GTPγS, CHAPSO and wildtype or E442Q spastin were mixed in this order to make a solution containing ∼60 µM tubulin, ∼60 µM spastin and 2 mM CHAPSO, and incubated at 30°C for 30 min. For E442Q spastin, 2 mM ATP was also added to this solution. Large aggregates were removed by centrifugation at 15,000 g, 30°C for 2 min, and 4 µL of the supernatant was placed on a glow-discharged holey carbon grid (R1.2/1.3; Quantifoil) inside the blotting machine kept at 30 °C and 100% humidity (Vitrobot IV; Thermo Fisher Scientific).

Grids were immediately blotted and plunged in liquid ethane. Data collection was performed using SerialEM software (*5*) on CRYO ARM 300 microscope (JEOL) equipped with a K3 camera (Gatan) in the electron counting mode, a cold-field emission gun, and an in-column Ω filter. Movies were acquired with the total dose of ∼60 e^-^/Å^2^ subdivided into 60 frames, 300 kV acceleration voltage, and target defocus value ranging −1.4 to −1.6 µm. 60,000× nominal magnification was used, which resulted in 0.75 Å/pixel.

Collected movies were analyzed using CryoSPARC (v4.7) as outlined in Figure S14. The movies were first processed by patch motion correction and patch CTF estimation. To make templates, ∼400 particles were manually picked and a 3D reconstruction was built with C4 symmetry. Particles were picked and selected by several rounds of template picking, 2D classification and heterogeneous refinement. 3D reconstruction was first built without imposing symmetry by *Ab-initio* reconstruction for the visual inspection to determine the symmetry (C14, C16 and C18) of each tubulin ring. To obtain high-resolution reconstruction of each tubulin ring, *Ab-initio* reconstruction jobs were conducted with C14, C16 and C18 symmetry, and masks for the corresponding rings were manually drawn using the 3D painting tool of napari or the volume eraser tool of ChimeraX. Each tubulin ring was then locally refined with these static masks. For refinement of the tubulin tetramer in the C16 ring, double stack particles were excluded, and the C14 ring and the C18 ring were subtracted using the reconstruction of the highest resolution. After C16 symmetry expansion, high-resolution volume was obtained by focused refinement of the tetramer. To improve map quality, particles were re-extracted, and refined by 3D classification and local refinement.

Atomic model of the tubulin tetramer in the C16 ring was built using Coot (v0.9.8.95). Initial atomic model was prepared by fitting α- and β-tubulin from deposited tubulin dimer model (PDB: 7YSO) to the refined map using ChimeraX “fitmap” command. Model refinement and validation were performed by Phenix (v1.21.2) (*6*) and the statistics are summarized in Table S1.

The structural parameters of tubulin tetramers (protofilament swing, dimer twist, and curvature) were calculated as follows. Atomic models listed in Table S2 were downloaded from RCSB Protein Data Bank, and their chain IDs were edited using the “changechains” command of ChimeraX so that their chain A, B, C and D correspond to the α-, β-, α- and β-tubulin from the minus to the plus end. For 3J2U and 7U0F, two atomic models were fitted to the corresponding density map (EMD-5565, EMD-26257) using the “fitmap” command of ChimeraX, because the atomic model only contain one set of αβ-tubulin. According to Figure S18, vectors ***x***_0_, ***y***_0_ and ***z***_0_ were calculated from chains A and B, while vectors ***x***_1_, ***y***_1_ and ***z***_1_ were calculated from chains C and D, using the center of mass of the indicated residues calculated by the Biopython package (*7*). Structural parameters in radians were defined as follows: protofilament swing = arctan(***ẑ***_1_ ⋅ ***x̂***_0_ /***ẑ***_1_ ⋅ ***ŷ***_0_), dimer twist = arctan(***ŷ***_1_ ⋅ ***x̂***_0_/***ŷ***_1_ ⋅ ***ŷ***_0_), curvature = arcsin(***ẑ***_0_ × ***ẑ***_1_ /|***ẑ***_0_ × ***ẑ***_1_ |), where the hat indicates normalization: ***v̂*** = ***v***/|***v***|.

Visualization of displacement vector maps and RMSD was done in ChimeraX. Each atomic model was fitted to the model of tubulin tetramer in the ring stack by the indicated chains using the “matchmaker” command. Displacement vectors of the corresponding Cα atoms in the β_2_ chain was calculated using a custom Python script and exported as text files containing arrow objects in the BILD format, which was directly loaded to ChimeraX. For the RMSD analysis, the ribbon-model object was colored by the RMSD values for each amino-acid residue that are available after running the “matchmaker” command.

## Data Analysis

All the statistical testing was conducted with Python 3.12.7, using scipy (*8*) or scikit-posthocs. Sample sizes are shown in the corresponding figure or legend. Details of the image analysis methods for TIRFM and electron microscopy are described in the corresponding section.

---

**Fig. S1.**
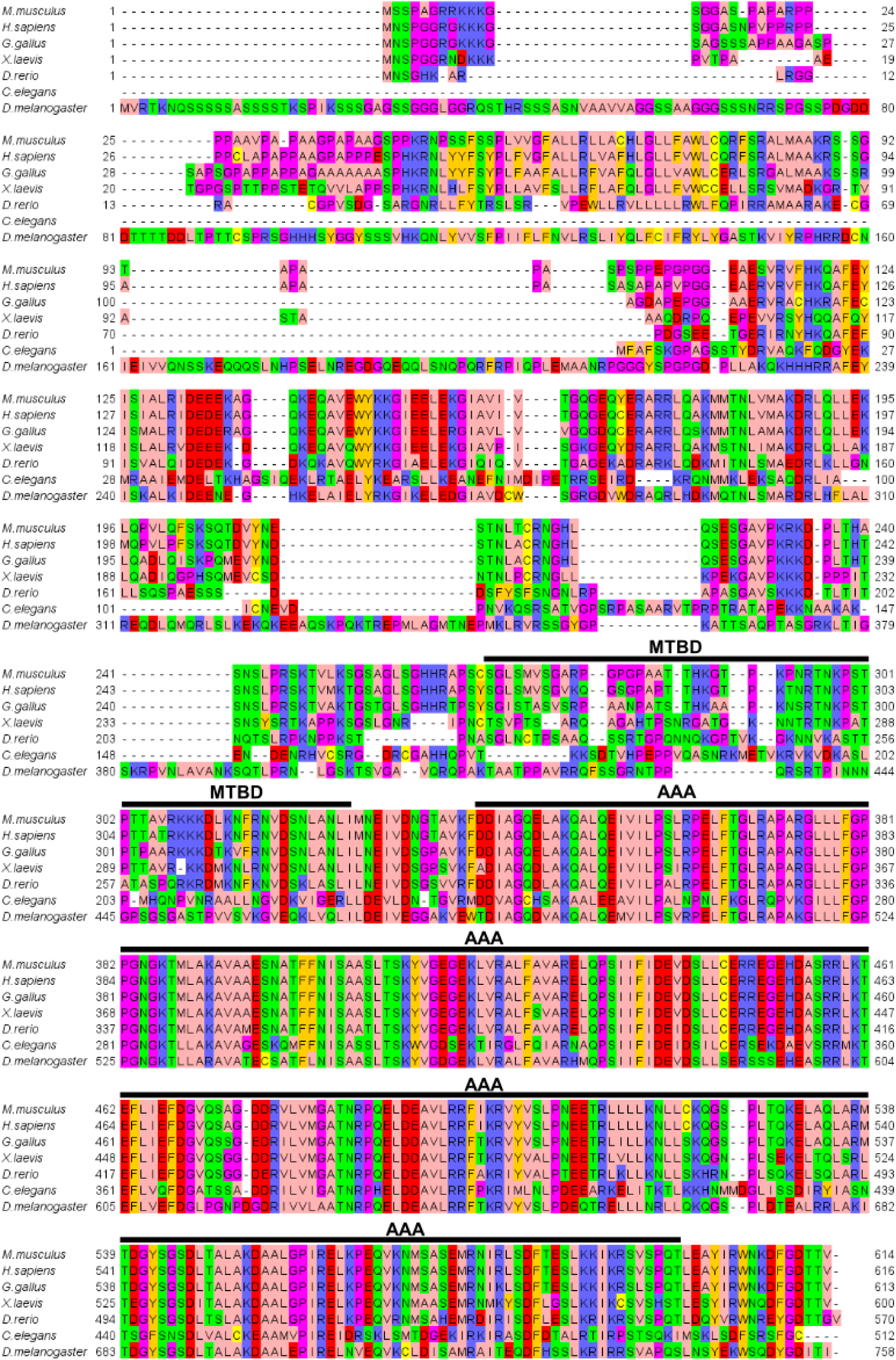
Alignment result of full-length spastin performed by JalView (v2.11.4.1).

**Fig. S2.**
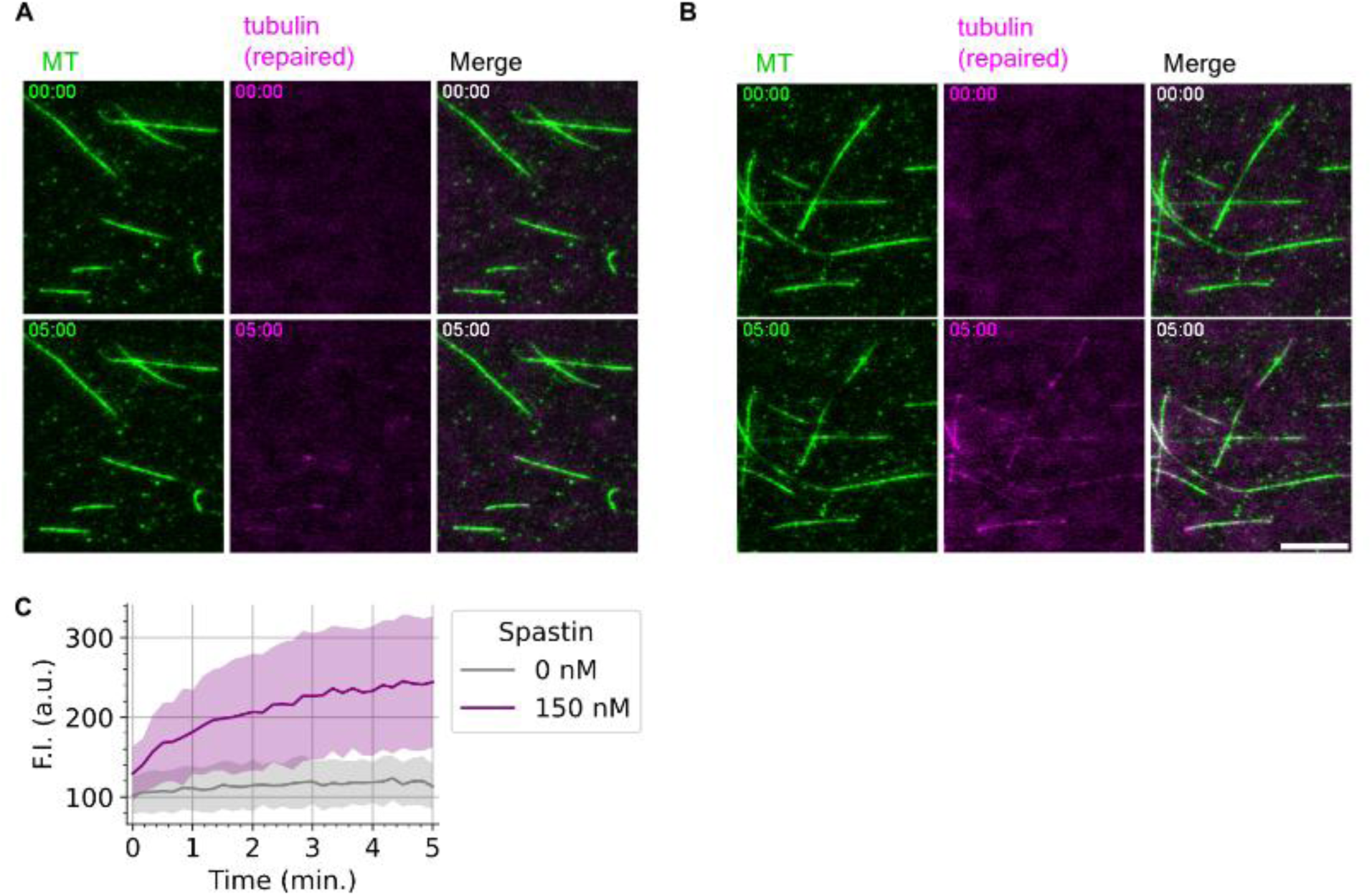
Microtubule lattice damage and repair assay. (**A**, **B**) Representative images showing incorporation of free tubulin without spastin (A), or with 150 nM spastin (B). Scale bar, 10 µm. (**C**) Quantification of the tubulin incorporation by the fluorescence intensity of free tubulin along microtubules. Lines and shadows indicate mean ± SD (one experiment; number of analyzed microtubules, 0 nM spastin: *n* = 28, 150 nM spastin: *n* = 32).

**Fig. S3.**
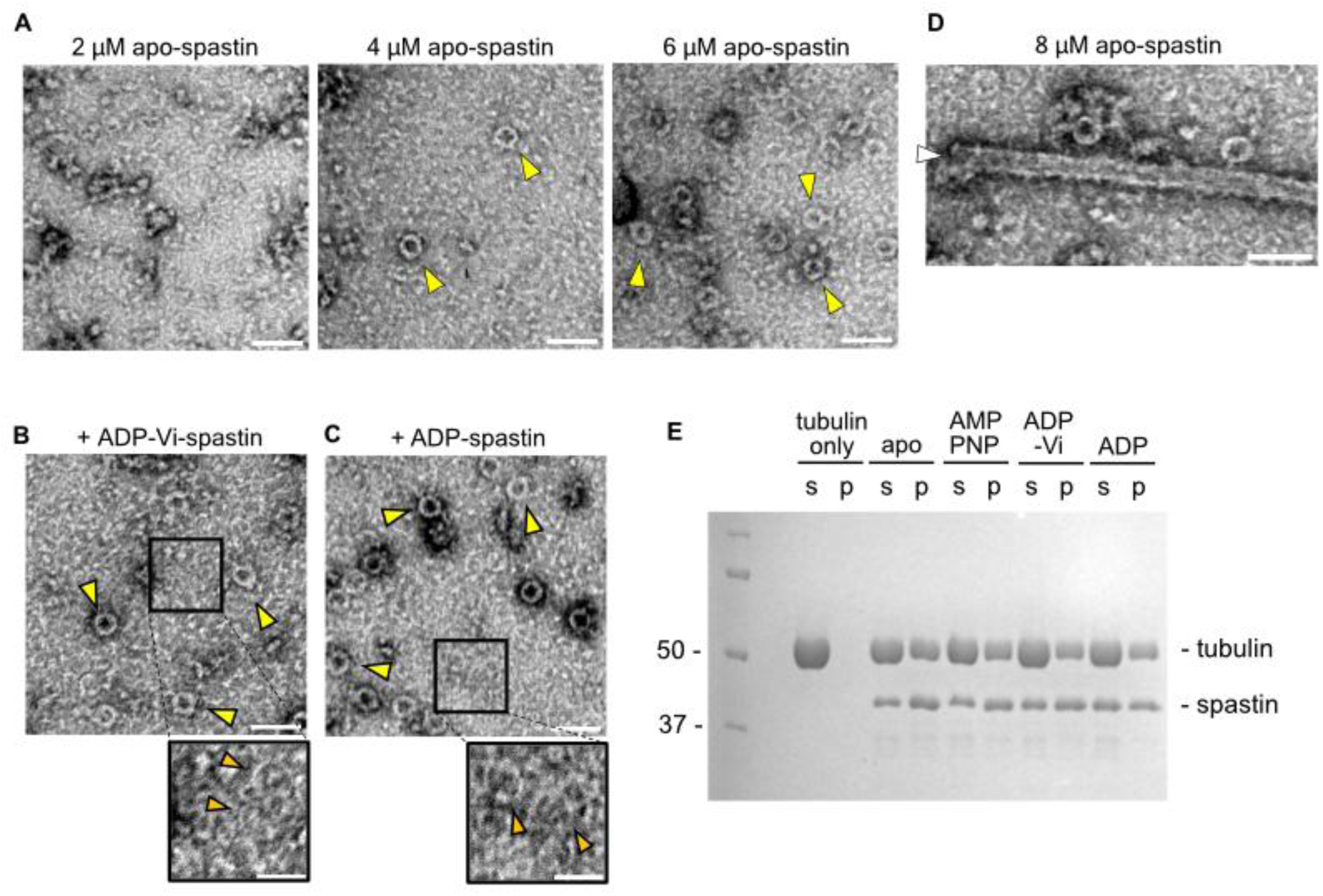
Tubulin ring formation by human spastin. (**A**) Negative-stain EM micrographs of mixture of 8 µM tubulin and different concentration of spastin. Arrowheads point at thick tubulin rings (yellow) and thin tubulin rings (orange). Scale bar, 100 nm. (**B**, **C**) Tubulin rings formed by spastin with ADP-Vi (B) or ADP (C), related to Fig. 1. Boxed regions are zoomed in the lower panels. Scale bar, 100 nm (upper) and 50 nm (lower). (**D**) Another position of the grid of Figure 1F, containing the mixture of 8 µM tubulin and 8 µM apo-spastin. Arrowhead points at a microtubule. Scale bar, 100 nm. (**E**) The SDS-PAGE result of the pelleting assay with different nucleotide. s: supernatant, p: pellet. In this pelleting assay, 8 µM tubulin and 8 µM spastin were mixed in 12.5 µL scale.

**Fig. S4.**
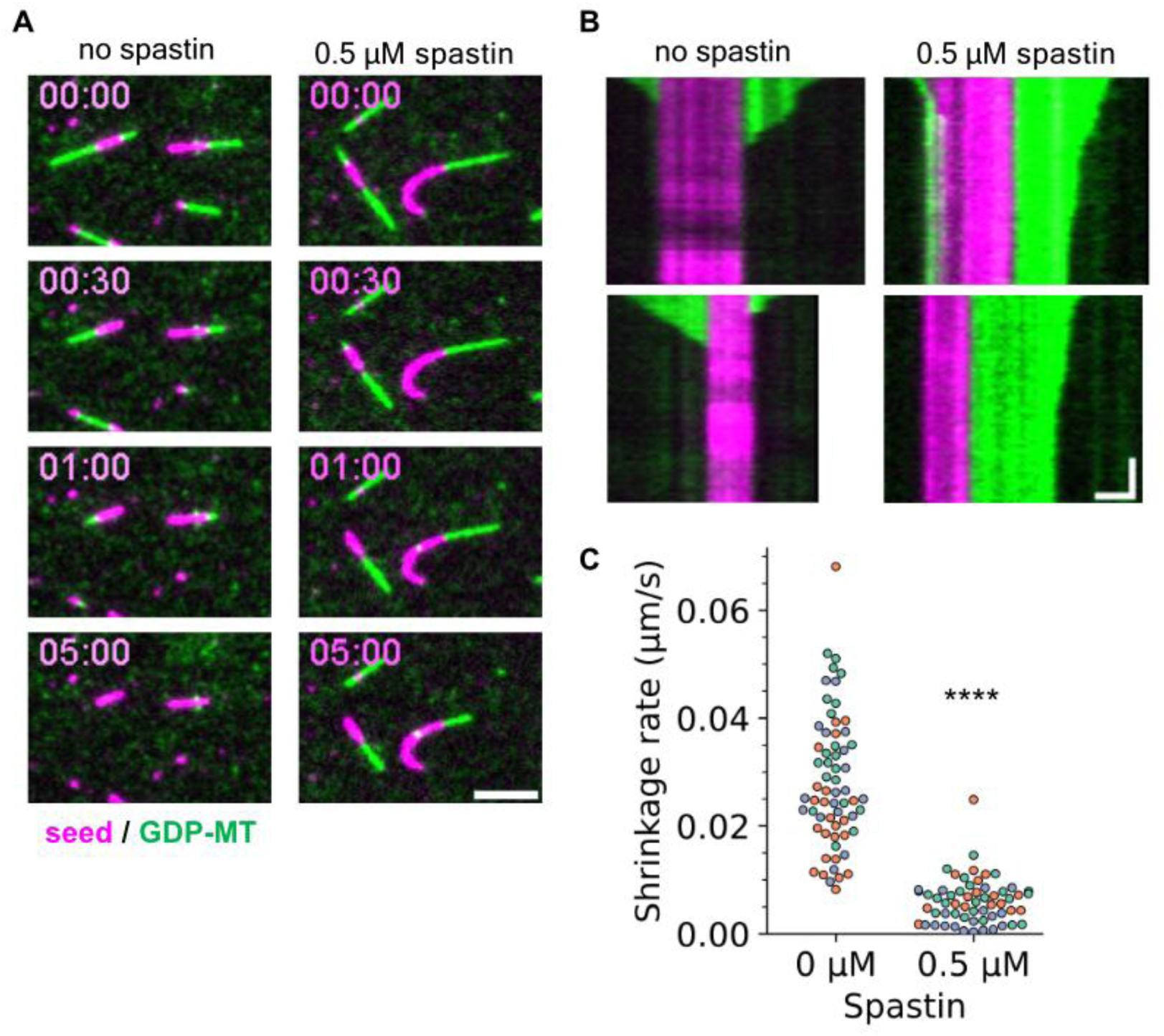
Depolymerization assay with human spastin. (**A**) Representative images showing spastin suppresses microtubule depolymerization. Scale bar, 5 µm. (**B**) Representative kymographs showing the shrinking microtubules. Scale bar, 2 µm (horizontal) and 1 min (vertical). (**C**) Plot of the microtubule shrinkage rate. Color represents each replicate of three experiments. ****: *p* < 0.0001 (Welch’s t-test).

**Fig. S5.**
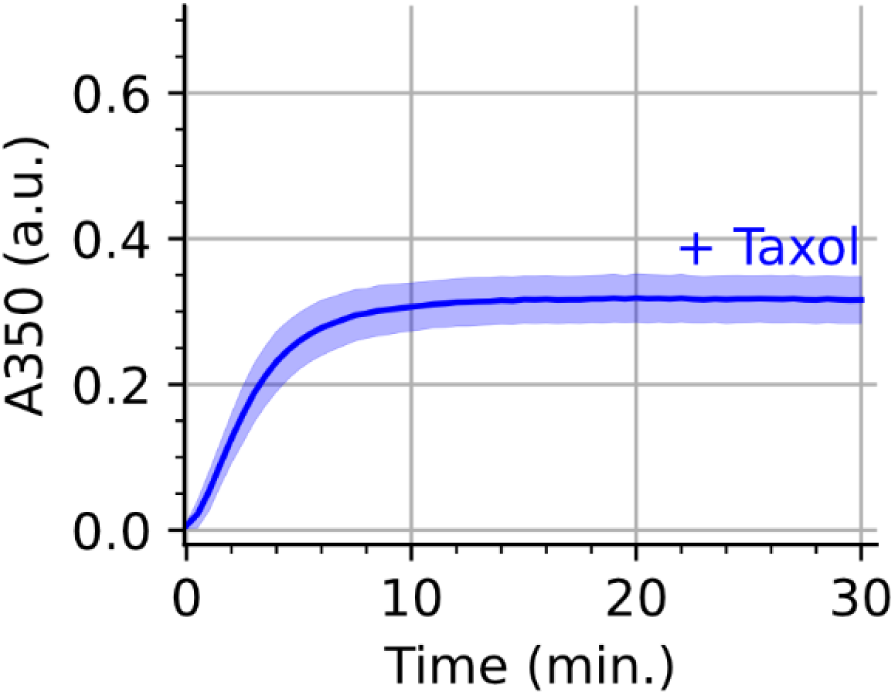
Turbidity assay with 10 µM taxol, related to エラー! 参照元が見つかりません。. **A.** Line and the shadow indicate mean ± SE (*n* = 3).

**Fig. S6.**
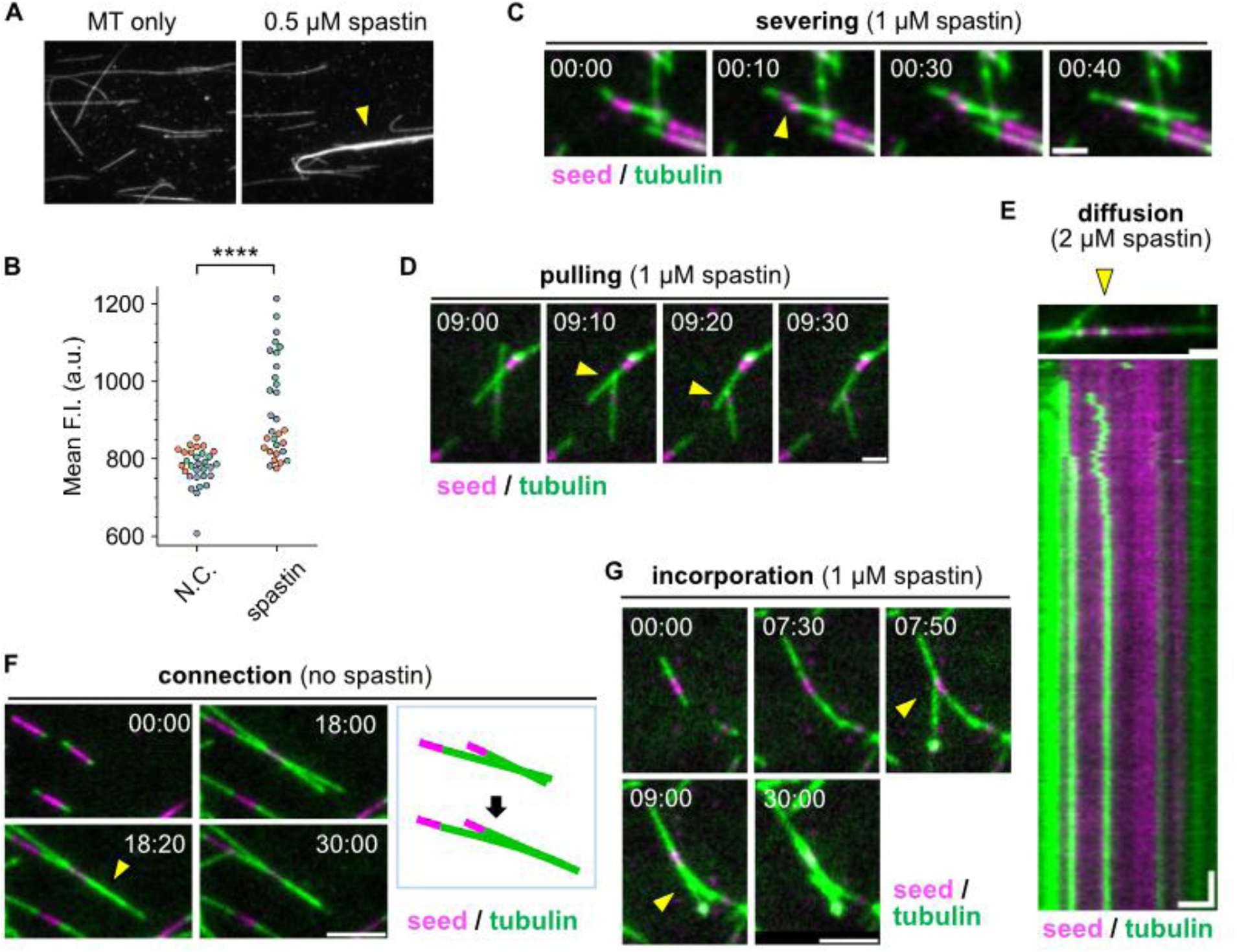
TIRF assays related to the microtubule bundling ability of spastin. (**A**) Representative images showing the spastin-dependent microtubule bundling (yellow arrowhead). (**B**) Quantification of the bundling efficiency by the mean fluorescence intensity on the microtubules for each shot. *n* = 30 image slices for each condition. Each color represents the replicate of three independent experiments. ****: *p* < 0.0001 (Welch’s t-test). (**C**) Representative image series of microtubule dynamics assay showing the severing event. The seed indicated by the arrowhead is severed, and immediately bundled to the original place. Scale bar, 2 µm. (**D**) Representative image series showing the pulling event. Arrowheads indicate the point where pulling force is generated. Scale bar, 2 µm. (**E**) A representative image and the corresponding kymograph of tubulin diffusively tethered on the seed. Scale bars, 2 µm (horizontal) and 2 min (vertical). (**F**) Representative image series of the bundling event without spastin. Arrowhead indicates the point where bundling occurred. Scale bar, 5 µm. (**G**) Representative image series showing an event that microtubule nucleated in the solution is incorporated into the microtubule elongated from a seed. Scale bar, 5 µm.

**Fig. S7.**
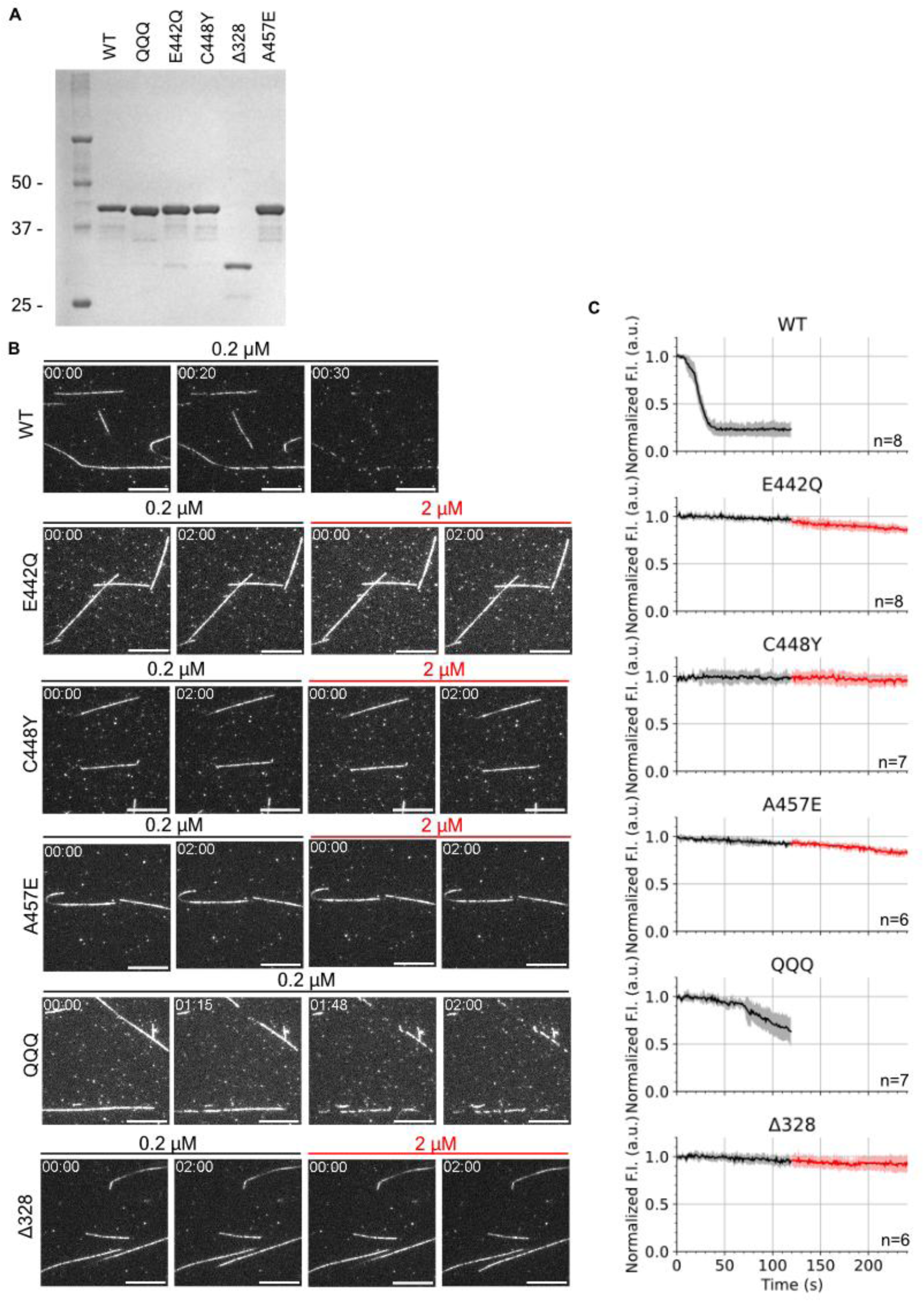
Severing activity of human spastin and the mutants. (**A**) SDS-PAGE of the purified proteins. 2 pmol was applied for each well. (**B**) Time-lapse images of the TIRFM-based severing assay for each mutant. Spastin concentration is shown above each image. Scale bar, 10 µm. (**C**) Quantification of severing by the fluorescence intensity of individual microtubules. Concentration was changed from 0.2 µM (black) to 2 µM (red) at the time point 120 s. Number of analyzed microtubules are shown in the plot (one experiment).

**Fig. S8.**
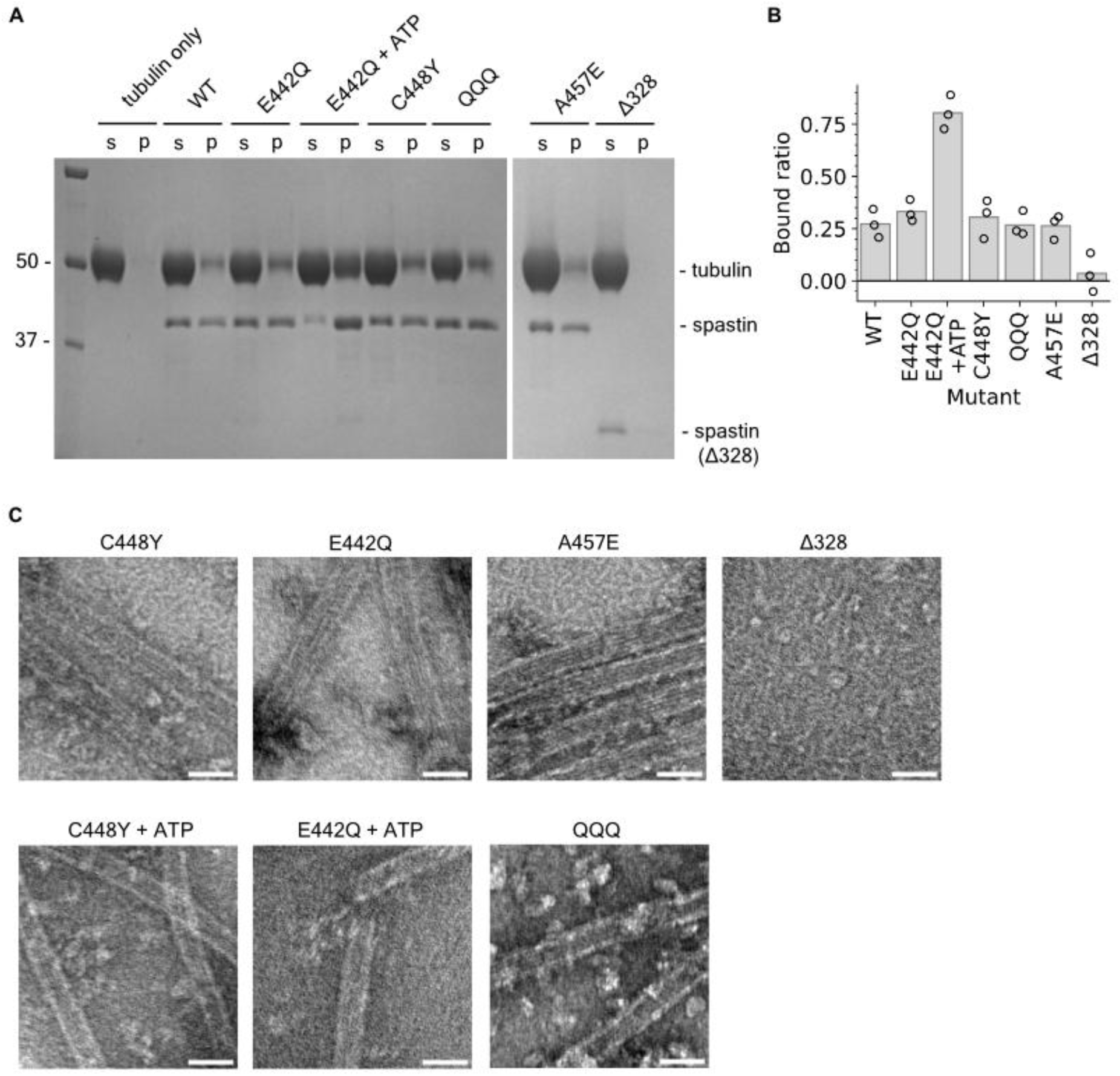
Characterization of the tubulin binding property of spastin mutants. (**A**) SDS-PAGE gels of the pelleting assay with spastin mutants. (**B**) Quantification of the bound ratio of spastin, calculated from the intensity of the spastin bands. (**C**) Negative-stain EM micrographs of microtubules formed after turbidity assays. Scale bar, 50 nm.

**Fig. S9.**
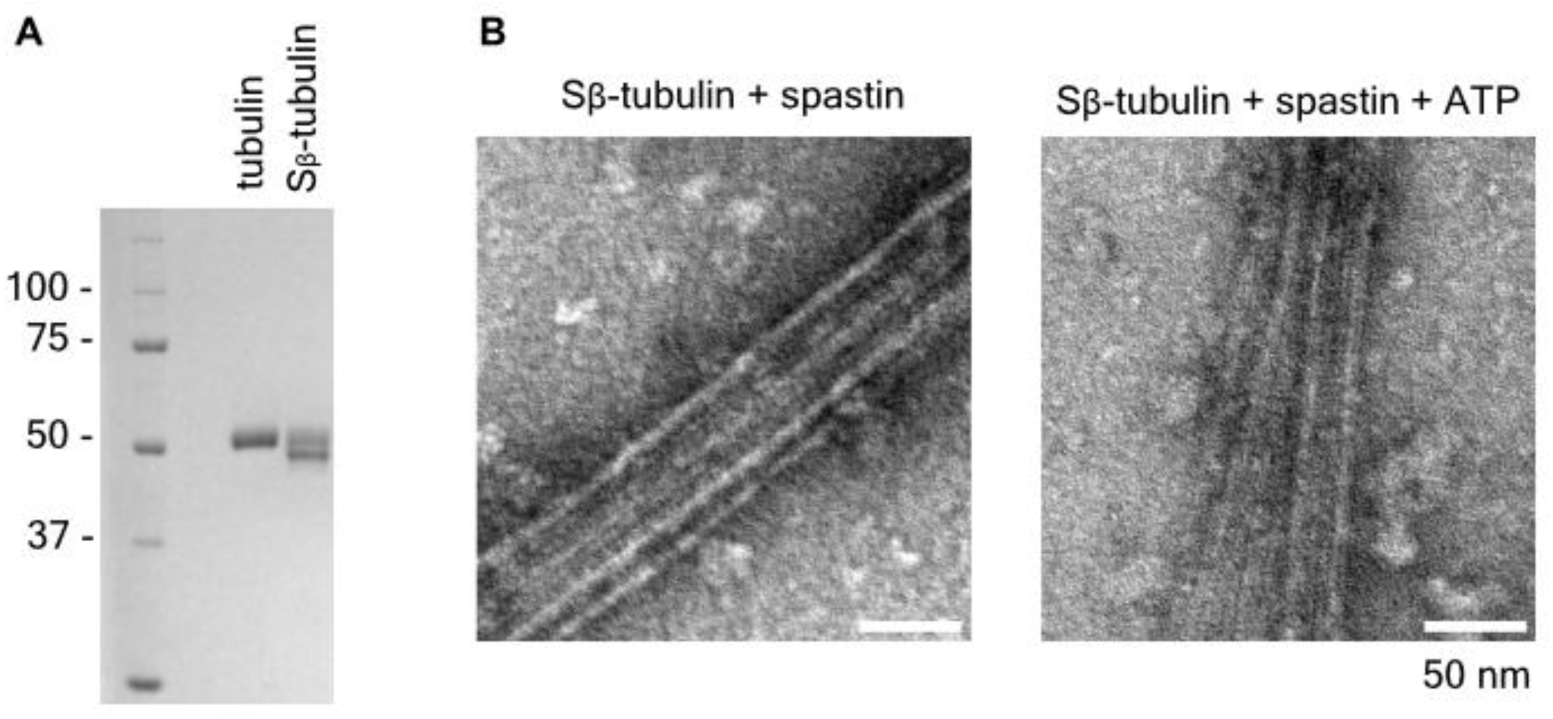
Nucleation of S_β_-tubulin by spastin. (**A**) SDS-PAGE of control brain tubulin and tubulin treated with subtilisin. β-CTT removal can be confirmed by the downward shift of the band. (**B**) Negative-stain micrographs of polymerized S_β_-tubulin after the turbidity assays.

**Fig. S10.**
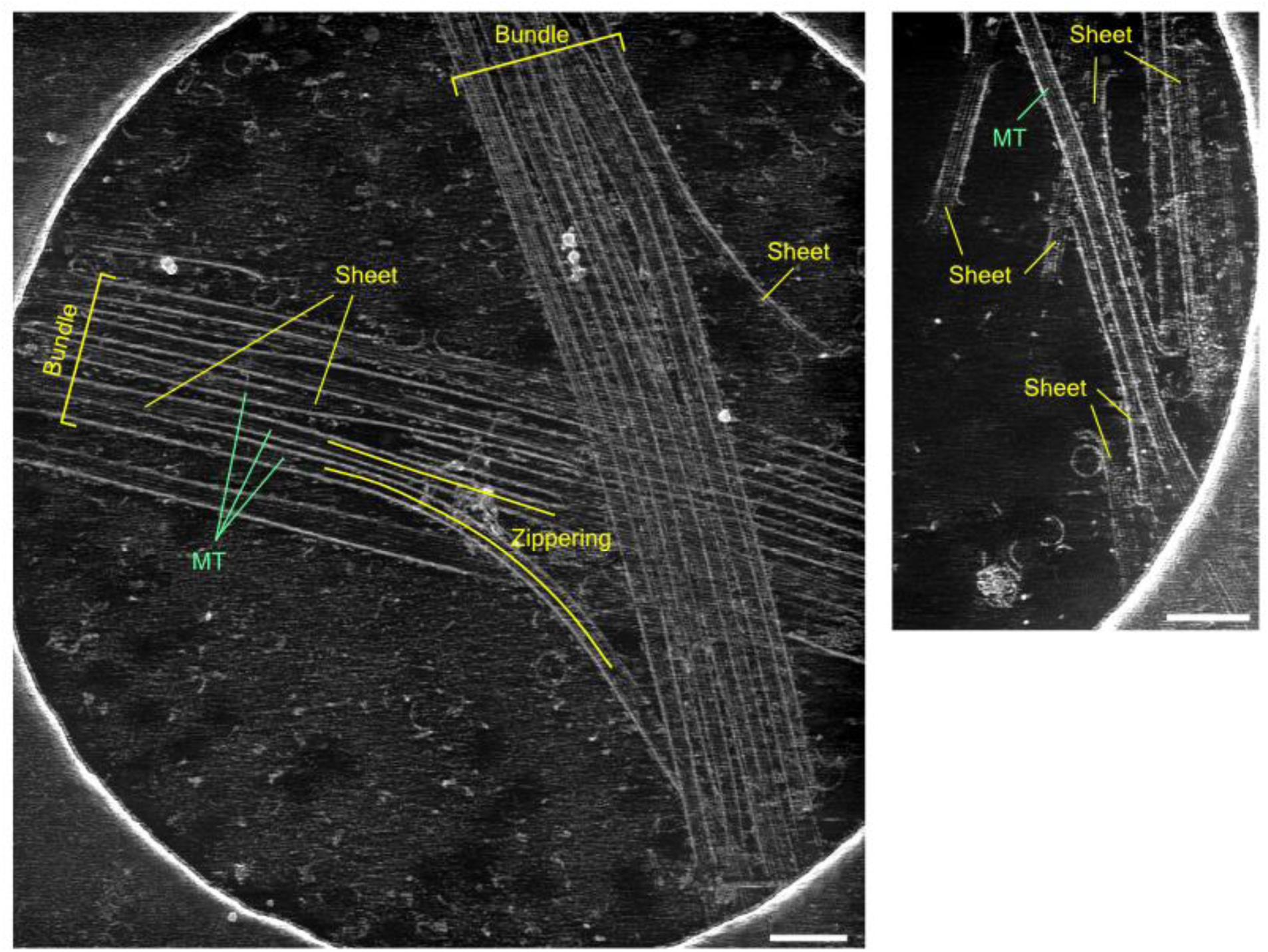
Max projection of reconstructed, denoised tomograms. Microtubule (MT), tubulin sheet, bundled microtubules and the zippering microtubule pair are labeled.

**Fig. S11.**
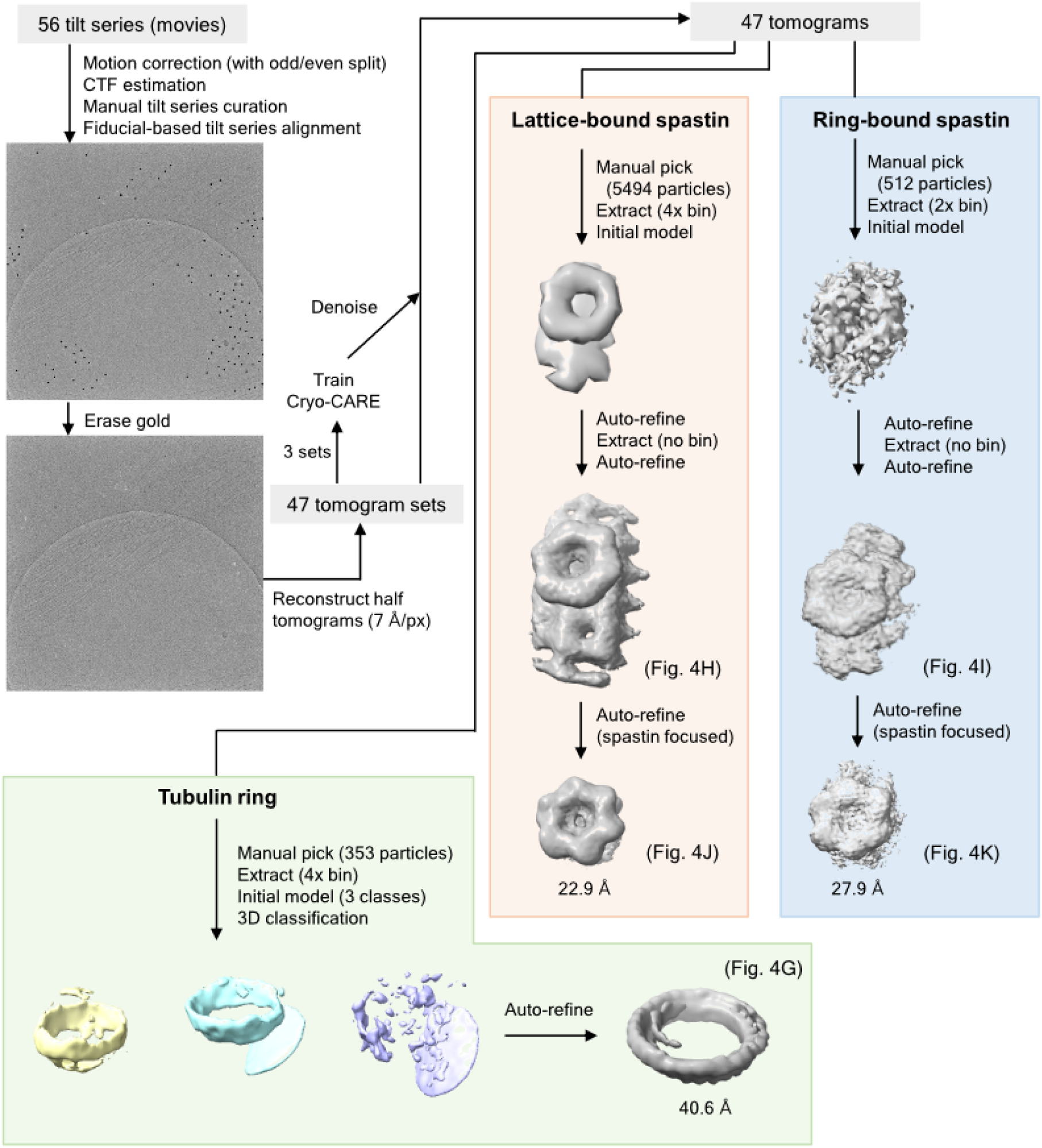
Outline of the analysis pipeline of cryo-ET. Resolutions shown next to the maps are the value at FSC=0.143.

**Fig. S12.**
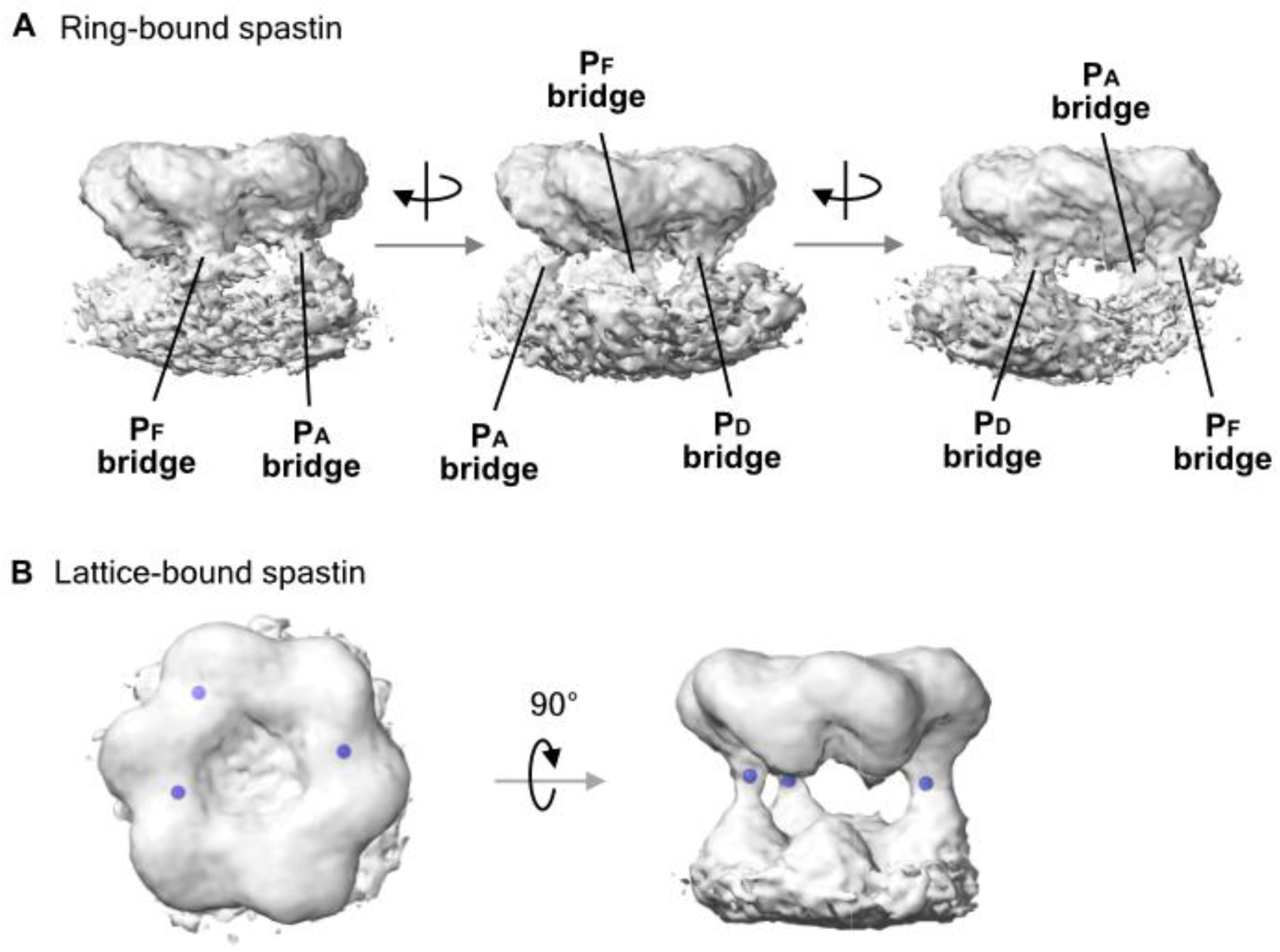
Details of the maps related to エラー! 参照元が見つかりません。4. (**A**) Spin view of the ring-bound spastin showing three bridges. (**B**) Raw ChimeraX snapshots of lattice-bound spastin with manually placed markers (blue) used for measuring distances.

**Fig. S13.**
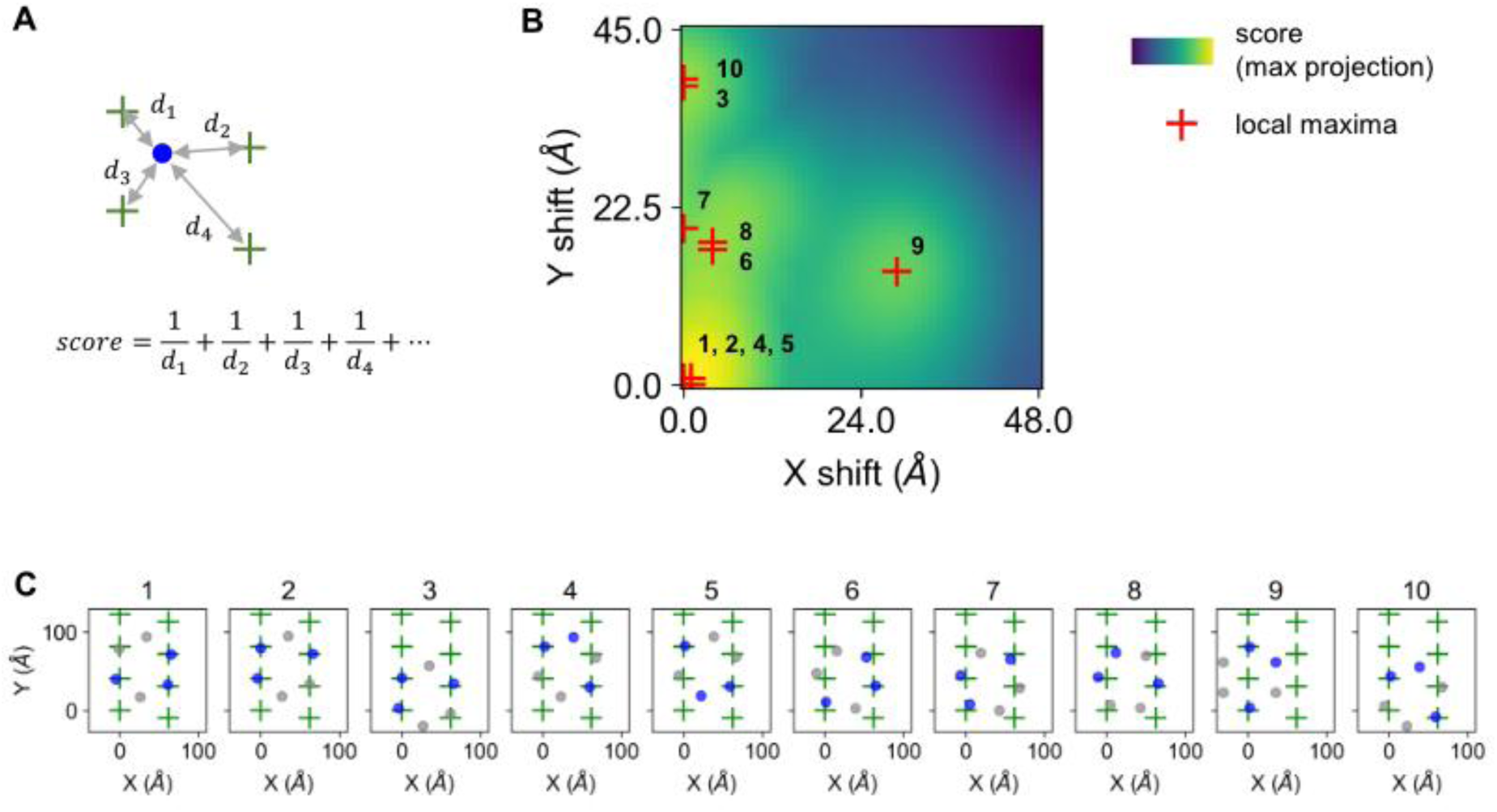
Exhaustive search of the possible poses of the spastin hexamer on the microtubule. (**A**) Schematic illustration of the model function of the score. Scores of all the combinations between eight tubulin CTTs and three spastin protomers with bridge (P_A_, P_D_, and P_F_) were calculated for every center position and rotation of the spastin hexamer. The X/Y shift indicates the shift from the center of the parallelogram of four tubulin monomers. (**B**) Maximum projection of the score. Red markers indicate the top-10 local maxima of the score. Numbers are the ranks. (**C**) Spastin poses with the top scores. Blue dots are the spastin protomers with bridge, for which scores are calculated. Numbers are the ranks as in (B).

**Fig. S14.**
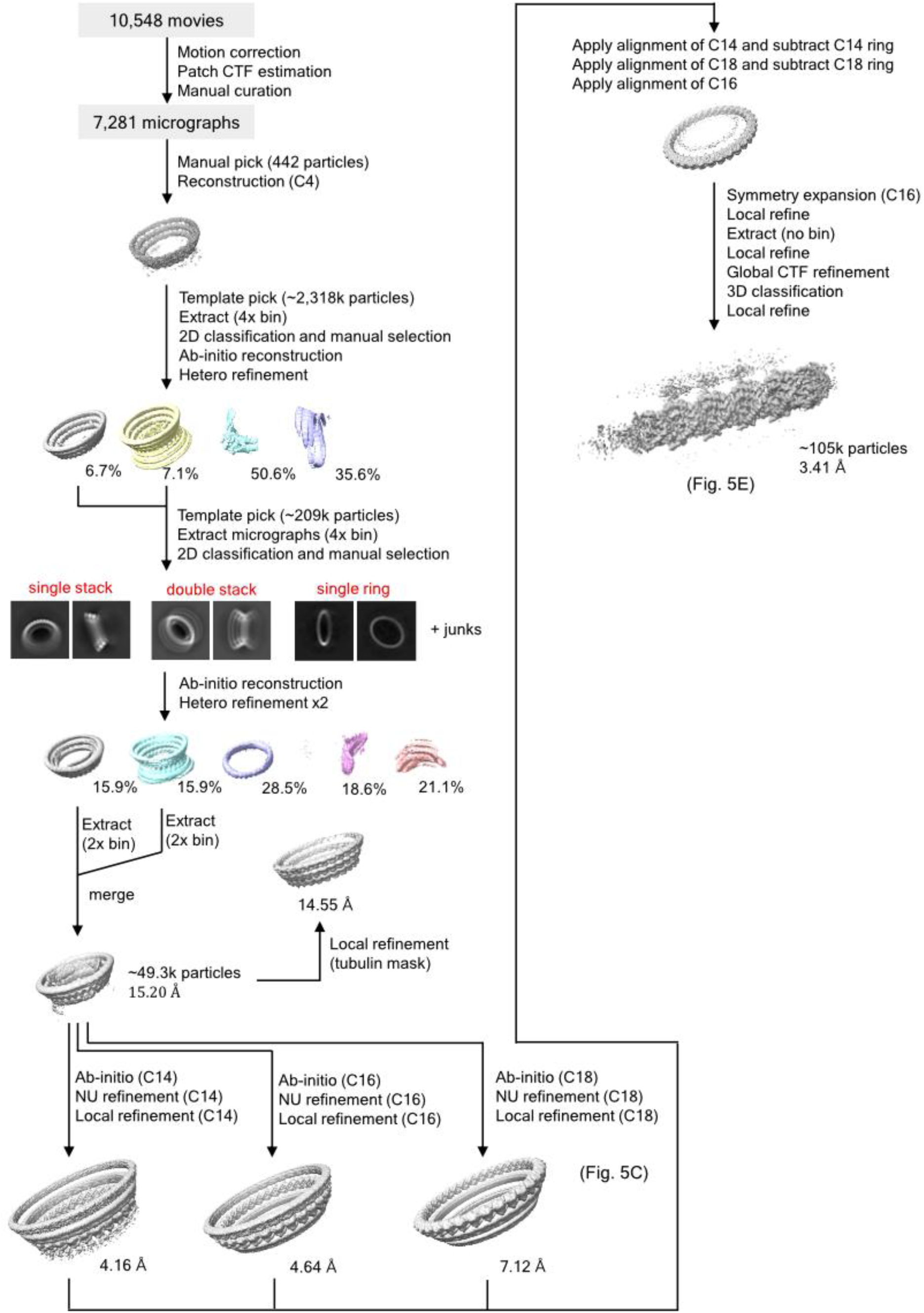
Outline of the single particle analysis of the tubulin ring stack formed by wildtype spastin. Resolutions shown next to the maps are the value at FSC=0.143.

**Fig. S15.**
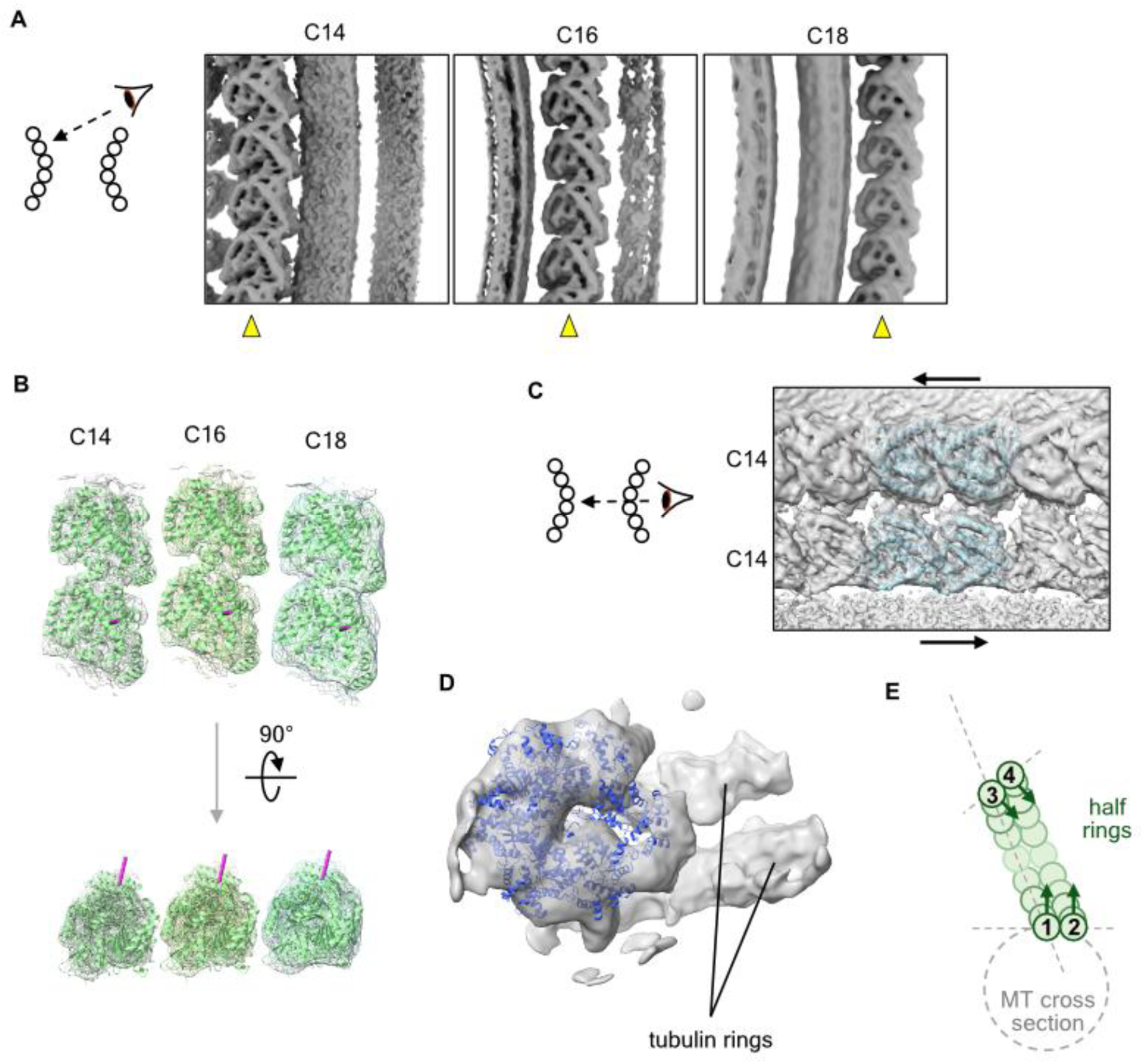
Details of the structure of tubulin rings. (**A**) Structures of tubulin rings. C14, C16 and C18 reconstructions are shown in separate panels. (**B**) Results of fitting tubulin dimers (PDB: 7YSO) to the C14, C16 and C18 reconstructions. Each magenta cylinder is the vector pointing from Ser170 to Met425 of the α-tubulin, which is almost parallel to the normal vector of the microtubule surface. (**C**) Zoom-in view of the C14-C14 interface showing that the protofilaments are antiparallel. (**D**) Density of the spastin hexamer inside the tubulin ring stack. An atomic model of the spastin hexamer (6PEN) is fitted. (**E**) Schematic illustration explaining the reason why stacking rings with increasing diameter is favorable. Green circles indicate tubulin monomers, and arrows on the sphere indicates the representative orientation of the tubulin that matches the normal vector of the microtubule surface. The interaction between tubulin 1 and 2 must be perpendicular to the arrows because of the favorable lateral contact of tubulin protofilaments, and so does the interaction between tubulin 3 and 4. The line connecting tubulin 1 and 3 is tilted against the arrow of tubulin 1 because of the favorable longitudinal contact of tubulin dimers.

**Fig. S16.**
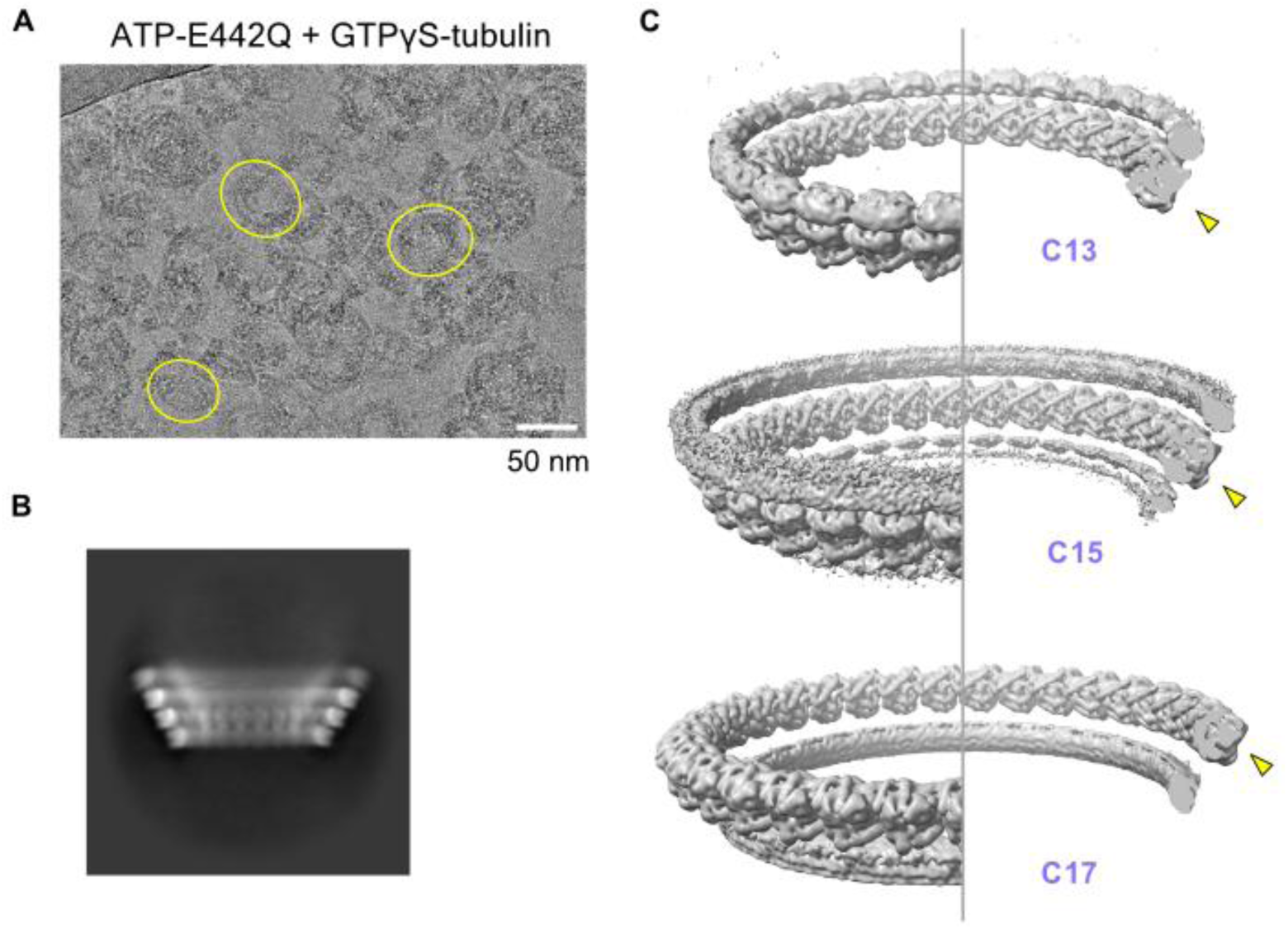
Cryo-EM data collection and the single-particle analysis of tubulin ring stack formed by E442Q spastin in the presence of ATP. (**A**) A representative, binned micrograph. Yellow circles indicate the tubulin ring stacks. (**B**) Projection of the C1 reconstruction of the tubulin ring stack induced by E442Q. (**C**) C13, C15 and C17 reconstruction. Arrowheads indicate the focused ring during refinement.

**Fig. S17.**
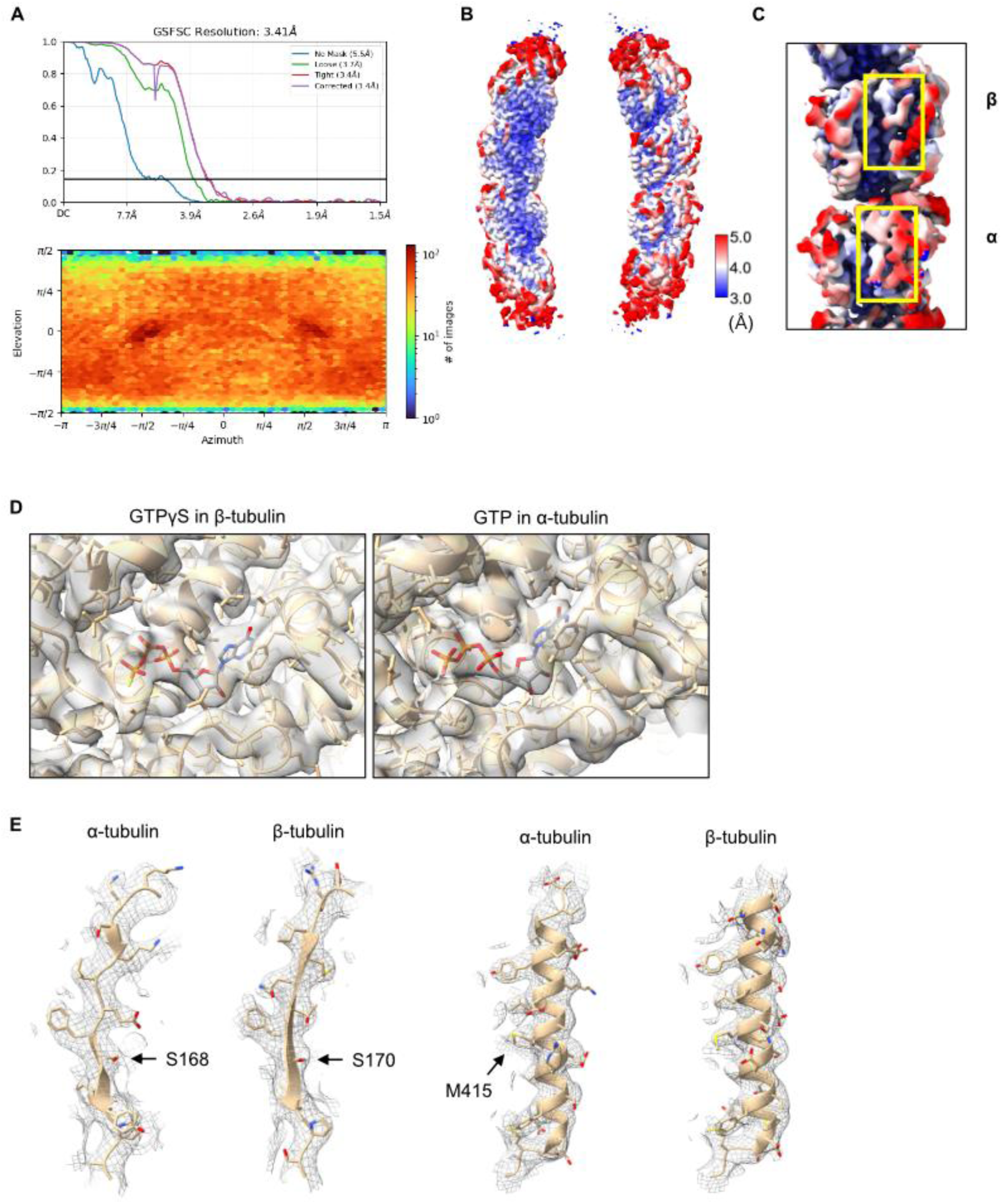
Validation of the reconstruction of the tubulin tetramer in the C16 tubulin ring. (**A**) Plot of the gold-standard Fourier shell correlation and the distribution of the particle orientation calculated in CryoSPARC. (**B**) Map of tubulin tetramer colored by the local-resolution estimation. (**C**) Zoom-in view of the tubulin tetramer highlighting the S9-S10 loop, a distinguishable feature between α- and β-tubulin. Map is colored by local-resolution as in (B). (**D**) Zoom-in view of the nucleotides. (**E**) Examples of tubulin secondary structures overlaid with the model side chains. The serine and methionine are the residues used for vector calculation (see also fig. S18B).

**Fig. S18.**
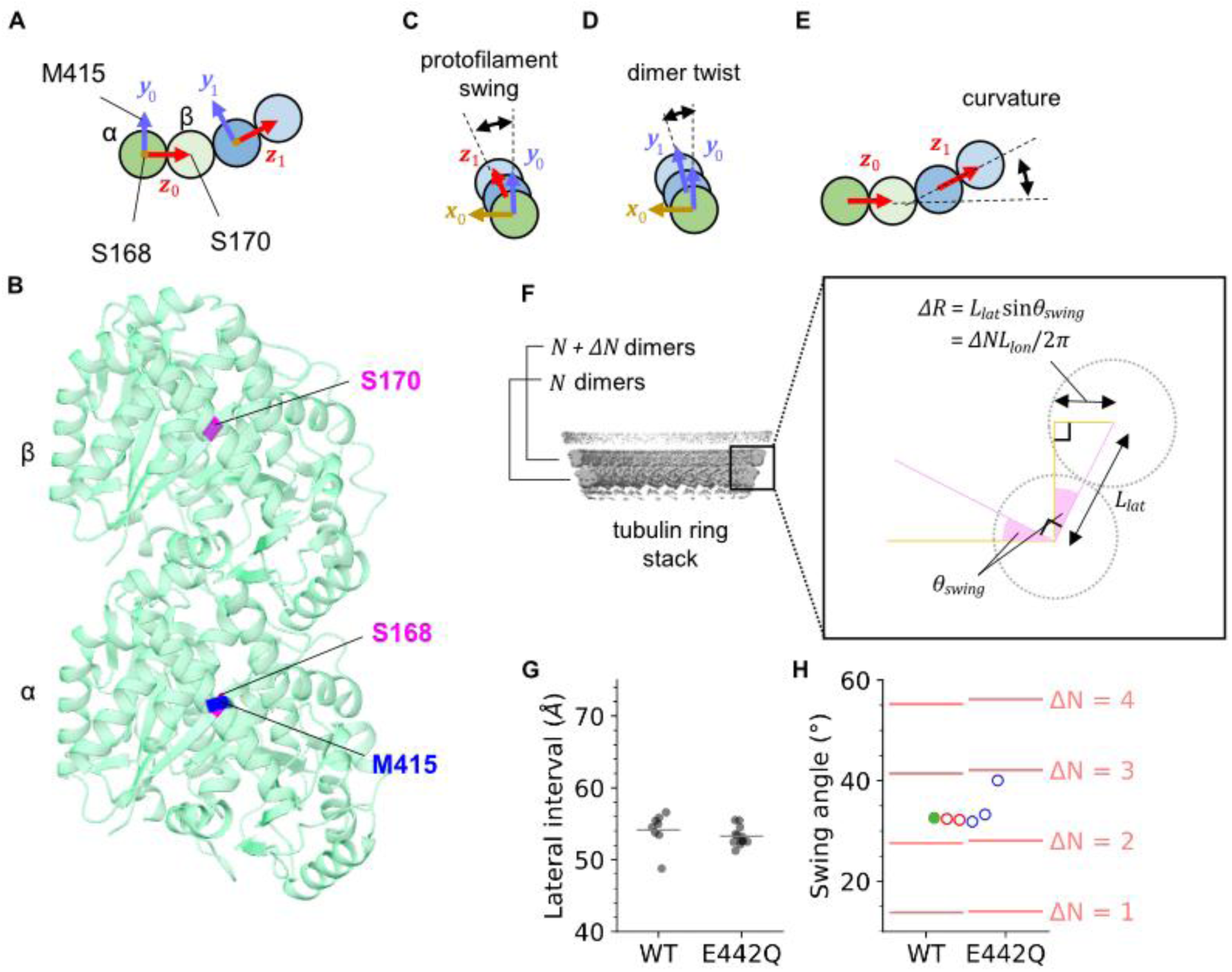
Calculation of structural parameters in the tubulin dimer interface. (**A**) Vectors defined by the atomic model of tubulin tetramers. ***x***_**0**_ and ***x***_**1**_ are calculated by ***x***_***i***_ = ***y***_***i***_ × ***z***_***i***_, and thus perpendicular to the plane of the paper. (**B**) Tubulin dimer viewed from outside of the microtubule. Residues used for calculation of vectors in (A) are colored. (**C-E**) Mathematical definition of the angles that represent the deformation of tubulin tetramers. Protofilament swing (C) and dimer twist (D) are calculated by projecting the vector in the next tubulin dimer to the ***x***_**0**_***y***_**0**_ plane, while curvature (E) is calculated by the rotation to match ***z***_**0**_ to ***z***_**1**_. (**F**) Theoretical prediction of the protofilament swing using the tubulin longitudinal and lateral intervals (𝑳_𝐥𝐨𝐧_, 𝑳_𝐥𝐚𝐭_), and 𝚫𝑵-dimer increment. This equation is derived by calculating the increment of tubulin ring radius 𝚫𝑹 in two ways: trigonometric ratio (𝚫𝑹 = 𝑳_𝒍𝒂𝒕_ 𝐬𝐢𝐧 𝜽_𝒔𝒘***i***𝒏𝒈_), and the difference in the perimeter of each tubulin ring (𝟐𝝅𝚫𝑹 = 𝚫𝑵𝑳_𝒍𝒐𝒏_). From this equation, protofilament swing is discretized as 𝜽_𝐬𝐰𝐢𝐧𝐠_ = 𝐚𝐫𝐜𝐬𝐢𝐧(𝚫𝑵𝑳_𝐥𝐨𝐧_/𝟐𝝅𝑳_𝐥𝐚𝐭_). (**G**) Plot of lateral interval (𝑳_𝐥𝐚𝐭_) measured from the C1 reconstruction of tubulin ring stacks formed by WT or E442Q spastin. The gray horizontal bars are the mean value of each dataset. (**H**) Plot of the theoretically achievable protofilament swing (𝜽_𝐬𝐰𝐢𝐧𝐠_) for each dimer increment number (𝚫𝑵), predicted using 𝑳_𝐥𝐨𝐧_ = 82 Å (*9*) and the measured 𝑳_𝐥𝐚𝐭_ in (G). Results from WT and E442Q indicate that the values of achievable 𝜽_𝐬𝐰𝐢𝐧𝐠_ are consistent between these conditions. Protofilament swing angles of the tetramers in Figure 6G are also plotted in the same color and order; left to right, ring stack (9X66), GDP-BeF_3_^-^ (6GZE), GDP-AlF_3_ (6S9E), stathmin (1FFX), stathmin colchcine (1SA0), and kinesin-13 (6BBN).

**Fig. S19.**
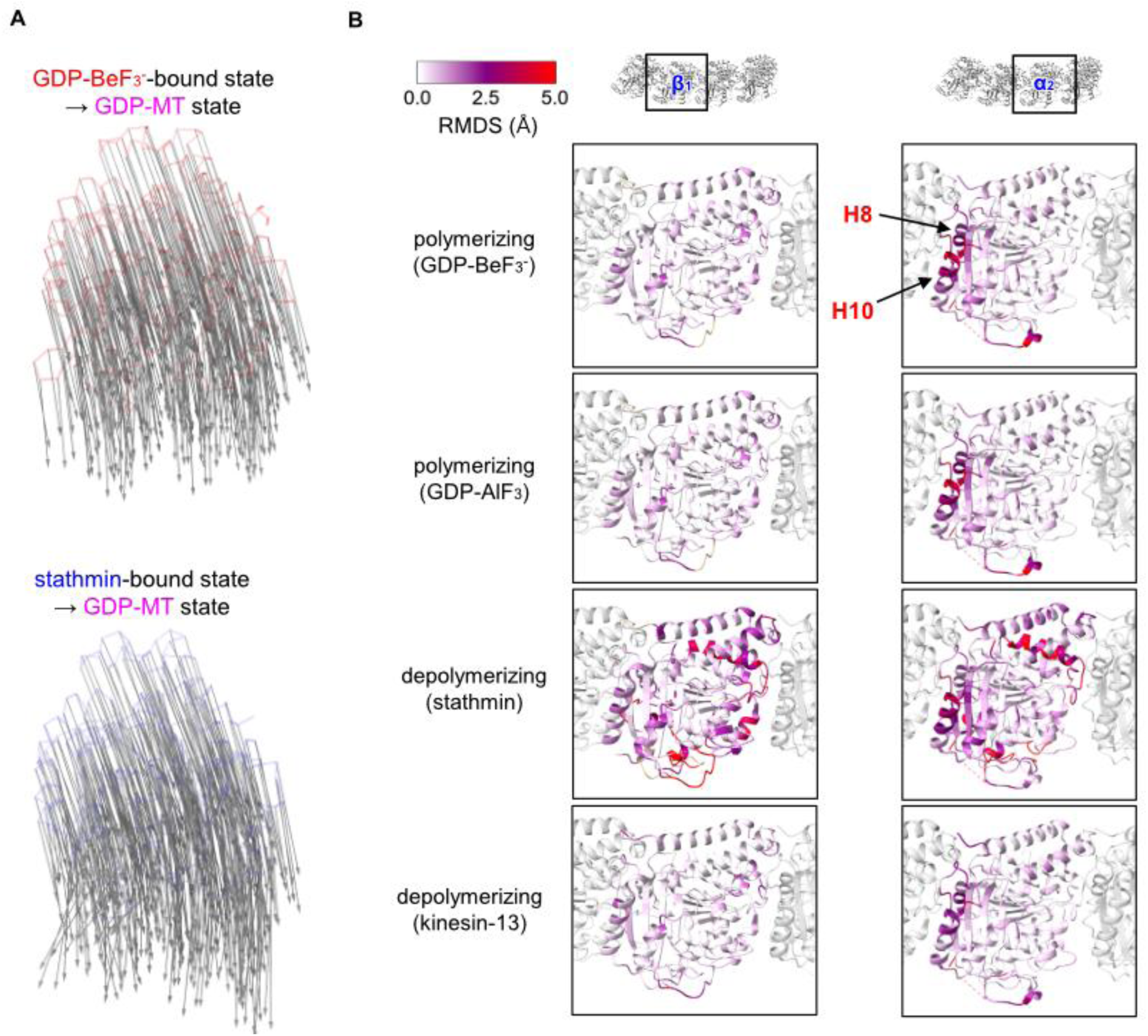
Analysis of conformational changes between tubulin tetramers. (**A**) Displacement vector maps related to Fig. 5H. (**B**) RMSD analysis of tubulin tetramers. Models were aligned on β_1_-tubulin to calculate β_1_ RMSD, and on α_2_-tubulin to calculate α_2_ RMSD. Each model is colored by the RMSD from the model of tubulin tetramer in the ring stack.

**Table S1.** Statistics of single particle analysis.

|  | **Tubulin tetramer in ring stack** |
| --- | --- |
| **Data Collection** |  |
| Microscope | CRYO ARM 300 |
| Camera | Gatan K3 |
| Magnification | 60,000 |
| Voltage (kV) | 300 |
| Electron dose (e<sup>-</sup>/Å<sup>2</sup>) | 60 |
| Defocus range (μm) | -1.6 – -1.4 μm |
| Pixel size (Å/pixel) | 0.75 |
| Particles | 104,987 |
| Symmetry | C1 |
| Resolution at FSC=0.143 (Å) | 3.41 |
| **Model composition** |  |
| Nonhydrogen atoms | 13631 |
| Protein residues | 1721 |
| Ligands | 2GTP, 2GTPγS |
| **R.m.s.deviation** |  |
| Bound lengths (Å) | 0.007 |
| Bond angles (°) | 0.932 |
| **Validation** |  |
| MolProbity Score | 1.84 |
| Clashscore | 12.11 |
| Rotamer outliers (%) | 0.95 |
| **Ramachandran plot** |  |
| Favored (%) | 96.30 |
| Allowed (%) | 3.64 |
| Outliers (%) | 0.06 |

**Table S2.** Swing angle and twist angle of tubulin tetramers.

|  | **PDB ID** | **Protofilament swing (°)** | **Dimer twist (°)** | **Curvature (°)** |
| --- | --- | --- | --- | --- |
| **Ring stack** | 9X66 | 32.6 | -0.80 | 22.3 |
| **GDP-BeF<sub>3</sub><sup>-</sup> tubulin** | 6GZE | 32.4 | 4.56 | 21.0 |
| **GDP-AlF<sub>3</sub> tubulin** | 6S9E | 32.2 | 4.53 | 20.0 |
| **Tubulin-stathmin** | 1FFX | 31.9 | 8.08 | 21.6 |
| **Tubulin-stathmin-colchicine** | 1SA0 | 33.3 | 9.46 | 21.5 |
| **Tubulin-kinesin-13** | 6BBN | 40.0 | 8.66 | 28.9 |
| **Tubulin-kinesin-13 tube** | 3J2U* | 9.8 | 3.21 | 23.8 |
| **Tubulin-Rev** | 7U0F* | 26.0 | 0.81 | 24.0 |
| **GDP-MT** | 6DPV | 0.2 | 0.48 | 0.0 |

## References and Notes

1. N. B. Gudimchuk, J. R. McIntosh, Regulation of microtubule dynamics, mechanics and function through the growing tip. Nat. Rev. Mol. Cell Biol. 22, 777–795 (2021).

2. A. Akhmanova, M. O. Steinmetz, Control of microtubule organization and dynamics: two ends in the limelight. Nat. Rev. Mol. Cell Biol. 16, 711–726 (2015).

3. A. W. Hunter, M. Caplow, D. L. Coy, W. O. Hancock, S. Diez, L. Wordeman, J. Howard, The kinesin-related protein MCAK is a microtubule depolymerase that forms an ATP- hydrolyzing complex at microtubule ends. Mol. Cell 11, 445–457 (2003).

4. M. Zanic, P. O. Widlund, A. A. Hyman, J. Howard, Synergy between XMAP215 and EB1 increases microtubule growth rates to physiological levels. Nat. Cell Biol. 15, 688–693 (2013).

5. L. Meißner, L. Niese, S. Diez, Helical motion and torque generation by microtubule motors. Curr. Opin. Cell Biol. 88, 102367 (2024).

6. S. Guha, A. Patil, H. Muralidharan, P. W. Baas, Mini-review: Microtubule sliding in neurons. Neurosci. Lett. 753, 135867 (2021).

7. S. R. Norris, S. Jung, P. Singh, C. E. Strothman, A. L. Erwin, M. D. Ohi, M. Zanic, R. Ohi, Microtubule minus-end aster organization is driven by processive HSET-tubulin clusters. Nat. Commun. 9, 2659 (2018).

8. T. Imasaki, S. Kikkawa, S. Niwa, Y. Saijo-Hamano, H. Shigematsu, K. Aoyama, K. Mitsuoka, T. Shimizu, M. Aoki, A. Sakamoto, Y. Tomabechi, N. Sakai, M. Shirouzu, S. Taguchi, Y. Yamagishi, T. Setsu, Y. Sakihama, E. Nitta, M. Takeichi, R. Nitta, CAMSAP2 organizes a γ-tubulin-independent microtubule nucleation centre through phase separation. Elife 11 (2022).

9. A. Roll-Mecak, F. J. McNally, Microtubule-severing enzymes. Curr. Opin. Cell Biol. 22, 96–103 (2010).

10. K. J. Evans, E. R. Gomes, S. M. Reisenweber, G. G. Gundersen, B. P. Lauring, Linking axonal degeneration to microtubule remodeling by Spastin-mediated microtubule severing. J. Cell Biol. 168, 599–606 (2005).

11. J. J. Hartman, R. D. Vale, Microtubule disassembly by ATP-dependent oligomerization of the AAA enzyme katanin. Science 286, 782–785 (1999).

12. S. Mukherjee, J. D. Diaz Valencia, S. Stewman, J. Metz, S. Monnier, U. Rath, A. B. Asenjo, R. A. Charafeddine, H. J. Sosa, J. L. Ross, A. Ma, D. J. Sharp, Human Fidgetin is a microtubule severing the enzyme and minus-end depolymerase that regulates mitosis. Cell Cycle 11, 2359–2366 (2012).

13. C. R. Sandate, A. Szyk, E. A. Zehr, G. C. Lander, A. Roll-Mecak, An allosteric network in spastin couples multiple activities required for microtubule severing. Nat. Struct. Mol. Biol. 26, 671–678 (2019).

14. H. Han, H. L. Schubert, J. McCullough, N. Monroe, M. D. Purdy, M. Yeager, W. I. Sundquist, C. P. Hill, Structure of spastin bound to a glutamate-rich peptide implies a hand- over-hand mechanism of substrate translocation. J. Biol. Chem. 295, 435–443 (2020).

15. E. Zehr, A. Szyk, G. Piszczek, E. Szczesna, X. Zuo, A. Roll-Mecak, Katanin spiral and ring structures shed light on power stroke for microtubule severing. Nat. Struct. Mol. Biol. 24, 717–725 (2017).

16. E. A. Zehr, A. Szyk, E. Szczesna, A. Roll-Mecak, Katanin grips the β-tubulin tail through an electropositive double spiral to sever microtubules. Dev. Cell 52, 118–131.e6 (2020).

17. M. Vietri, K. O. Schink, C. Campsteijn, C. S. Wegner, S. W. Schultz, L. Christ, S. B. Thoresen, A. Brech, C. Raiborg, H. Stenmark, Spastin and ESCRT-III coordinate mitotic spindle disassembly and nuclear envelope sealing. Nature 522, 231–235 (2015).

18. F. J. McNally, A. Roll-Mecak, Microtubule-severing enzymes: From cellular functions to molecular mechanism. J. Cell Biol. 217, 4057–4069 (2018).

19. J. C. M. Meiring, I. Grigoriev, W. Nijenhuis, L. C. Kapitein, A. Akhmanova, Opto-katanin, an optogenetic tool for localized, microtubule disassembly. Curr. Biol. 32, 4660–4674.e6 (2022).

20. G. Y. Liu, S.-C. Chen, G.-H. Lee, K. Shaiv, P.-Y. Chen, H. Cheng, S.-R. Hong, W.-T. Yang, S.-H. Huang, Y.-C. Chang, H.-C. Wang, C.-L. Kao, P.-C. Sun, M.-H. Chao, Y.-Y. Lee, M.-J. Tang, Y.-C. Lin, Precise control of microtubule disassembly in living cells. EMBO J. 41, e110472 (2022).

21. W. Yu, L. Qiang, J. M. Solowska, A. Karabay, S. Korulu, P. W. Baas, The microtubule- severing proteins spastin and katanin participate differently in the formation of axonal branches. Mol. Biol. Cell 19, 1485–1498 (2008).

22. Z. Ji, G. Zhang, L. Chen, J. Li, Y. Yang, C. Cha, J. Zhang, H. Lin, G. Guo, Spastin interacts with CRMP5 to promote neurite outgrowth by controlling the microtubule dynamics: Spastin interacts with CRMP5 to promote neurite outgrowth. Dev. Neurobiol. 78, 1191– 1205 (2018).

23. A. Ma, Z. Liang, H. Zhang, Z. Meng, J. Zhu, S. Chen, Q. Lin, T. Jiang, M. Tan, UCHL1- mediated spastin degradation regulates microtubule severing and hippocampal neurite outgrowth. J. Mol. Neurosci. 75, 54 (2025).

24. T. Advedissian, S. Frémont, A. Echard, Cytokinetic abscission requires actin-dependent microtubule severing. Nat. Commun. 15, 1949 (2024).

25. J. Guizetti, L. Schermelleh, J. Mäntler, S. Maar, I. Poser, H. Leonhardt, T. Müller-Reichert, D. W. Gerlich, Cortical constriction during abscission involves helices of ESCRT-III- dependent filaments. Science 331, 1616–1620 (2011).

26. J. M. Solowska, P. W. Baas, Hereditary spastic paraplegia SPG4: what is known and not known about the disease. Brain 138, 2471–2484 (2015).

27. A. T. Lopes, T. J. Hausrat, F. F. Heisler, K. V. Gromova, F. L. Lombino, T. Fischer, L. Ruschkies, P. Breiden, E. Thies, I. Hermans-Borgmeyer, M. Schweizer, J. R. Schwarz, C. Lohr, M. Kneussel, Spastin depletion increases tubulin polyglutamylation and impairs kinesin-mediated neuronal transport, leading to working and associative memory deficits. PLoS Biol. 18, e3000820 (2020).

28. J. D. Wood, J. A. Landers, M. Bingley, C. J. McDermott, V. Thomas-McArthur, L. J. Gleadall, P. J. Shaw, V. T. Cunliffe, The microtubule-severing protein Spastin is essential for axon outgrowth in the zebrafish embryo. Hum. Mol. Genet. 15, 2763–2771 (2006).

29. L. Qiang, E. Piermarini, H. Muralidharan, W. Yu, L. Leo, L. E. Hennessy, S. Fernandes, T. Connors, P. L. Yates, M. Swift, L. V. Zholudeva, M. A. Lane, G. Morfini, G. M. Alexander, T. D. Heiman-Patterson, P. W. Baas, Hereditary spastic paraplegia: gain-of-function mechanisms revealed by new transgenic mouse. Hum. Mol. Genet. 28, 1136–1152 (2019).

30. A. Roll-Mecak, R. D. Vale, Making more microtubules by severing: a common theme of noncentrosomal microtubule arrays?, The journal of cell biology. 175 (2006)pp. 849–851.

31. A. Vemu, E. Szczesna, E. A. Zehr, J. O. Spector, N. Grigorieff, A. M. Deaconescu, A. Roll- Mecak, Severing enzymes amplify microtubule arrays through lattice GTP-tubulin incorporation. Science 361 (2018).

32. H. de Forges, A. Pilon, I. Cantaloube, A. Pallandre, A.-M. Haghiri-Gosnet, F. Perez, C. Poüs, Localized mechanical stress promotes microtubule rescue. Curr. Biol. 26, 3399–3406 (2016).

33. A. Aher, D. Rai, L. Schaedel, J. Gaillard, K. John, Q. Liu, M. Altelaar, L. Blanchoin, M. Thery, A. Akhmanova, CLASP mediates microtubule repair by restricting lattice damage and regulating tubulin incorporation. Curr. Biol. 30, 2175–2183.e6 (2020).

34. Y.-W. Kuo, O. Trottier, M. Mahamdeh, J. Howard, Spastin is a dual-function enzyme that severs microtubules and promotes their regrowth to increase the number and mass of microtubules. Proceedings of the National Academy of Sciences 116, 5533–5541 (2019).

35. M. L. Valenstein, A. Roll-Mecak, Graded control of microtubule severing by tubulin glutamylation. Cell 164, 911–921 (2016).

36. A. Pisciottani, L. Biancolillo, M. Ferrara, D. Valente, F. Sardina, L. Monteonofrio, S. Camerini, M. Crescenzi, S. Soddu, C. Rinaldo, HIPK2 phosphorylates the microtubule- severing enzyme spastin at S268 for abscission. Cells 8, 684 (2019).

37. R. Tan, A. J. Lam, T. Tan, J. Han, D. W. Nowakowski, M. Vershinin, S. Simó, K. M. Ori-McKenney, R. J. McKenney, Microtubules gate tau condensation to spatially regulate microtubule functions. Nat. Cell Biol. 21, 1078–1085 (2019).

38. A. Roll-Mecak, R. D. Vale, Structural basis of microtubule severing by the hereditary spastic paraplegia protein spastin. Nature 451, 363–367 (2008).

39. S. R. White, K. J. Evans, J. Lary, J. L. Cole, B. Lauring, Recognition of C-terminal amino acids in tubulin by pore loops in Spastin is important for microtubule severing. J. Cell Biol. 176, 995–1005 (2007).

40. T. Eckert, D. T.-V. Le, S. Link, L. Friedmann, G. Woehlke, Spastin’s microtubule-binding properties and comparison to katanin. PLoS One 7, e50161 (2012).

41. S. C. Shin, S.-K. Im, E.-H. Jang, K. S. Jin, E.-M. Hur, E. E. Kim, Structural and Molecular Basis for Katanin-Mediated Severing of Glutamylated Microtubules. Cell Rep. 26, 1357–1367.e5 (2019).

42. M. E. Bailey, D. L. Sackett, J. L. Ross, Katanin severing and binding microtubules are inhibited by tubulin carboxy tails. Biophys. J. 109, 2546–2561 (2015).

43. E. M. Mandelkow, E. Mandelkow, R. A. Milligan, Microtubule dynamics and microtubule caps: a time-resolved cryo-electron microscopy study. J. Cell Biol. 114, 977–991 (1991).

44. J. R. McIntosh, E. O’Toole, G. Morgan, J. Austin, E. Ulyanov, F. Ataullakhanov, N. Gudimchuk, Microtubules grow by the addition of bent guanosine triphosphate tubulin to the tips of curved protofilaments. J. Cell Biol. 217, 2691–2708 (2018).

45. D. Tan, A. B. Asenjo, V. Mennella, D. J. Sharp, H. Sosa, Kinesin-13s form rings around microtubules. J. Cell Biol. 175, 25–31 (2006).

46. R. Melki, M. F. Carlier, D. Pantaloni, S. N. Timasheff, Cold depolymerization of microtubules to double rings: geometric stabilization of assemblies. Biochemistry 28, 9143– 9152 (1989).

47. M. W. Kirschner, R. C. Williams, The mechanism of microtubule assembly in vitro. J. Supramol. Struct. 2, 412–428 (1974).

48. D. Portran, L. Schaedel, Z. Xu, M. Théry, M. V. Nachury, Tubulin acetylation protects long-lived microtubules against mechanical ageing. Nat. Cell Biol. 19, 391–398 (2017).

49. I. Arnal, C. Heichette, G. S. Diamantopoulos, D. Chrétien, CLIP-170/tubulin-curved oligomers coassemble at microtubule ends and promote rescues. Curr. Biol. 14, 2086–2095 (2004).

50. S. W. Manka, C. A. Moores, The role of tubulin-tubulin lattice contacts in the mechanism of microtubule dynamic instability. Nat. Struct. Mol. Biol. 25, 607–615 (2018).

51. J. An, T. Imasaki, A. Narita, S. Niwa, R. Sasaki, T. Makino, R. Nitta, M. Kikkawa, Dimerization of GAS2 mediates crosslinking of microtubules and F-actin. EMBO J., doi: 10.1038/s44318-025-00415-2 (2025).

52. F. Gaskin, C. R. Cantor, M. L. Shelanski, Turbidimetric studies of the in vitro assembly and disassembly of porcine neurotubules. J. Mol. Biol. 89, 737–755 (1974).

53. A. Thawani, M. J. Rale, N. Coudray, G. Bhabha, H. A. Stone, J. W. Shaevitz, S. Petry, The transition state and regulation of γ-TuRC-mediated microtubule nucleation revealed by single molecule microscopy. Elife 9 (2020).

54. M. Wieczorek, S. Bechstedt, S. Chaaban, G. J. Brouhard, Microtubule-associated proteins control the kinetics of microtubule nucleation. Nat. Cell Biol. 17, 907–916 (2015).

55. S. Salinas, R. E. Carazo-Salas, C. Proukakis, J. M. Cooper, A. E. Weston, G. Schiavo, T. T. Warner, Human spastin has multiple microtubule-related functions: Bundling and severing activities of human spastin. J. Neurochem. 95, 1411–1420 (2005).

56. V. Campanacci, A. Urvoas, S. Cantos-Fernandes, M. Aumont-Nicaise, A.-A. Arteni, C. Velours, M. Valerio-Lepiniec, B. Dreier, A. Plückthun, A. Pilon, C. Poüs, P. Minard, B. Gigant, Insight into microtubule nucleation from tubulin-capping proteins. Proc. Natl. Acad. Sci. U. S. A. 116, 9859–9864 (2019).

57. H. Drechsler, Y. Xu, V. F. Geyer, Y. Zhang, S. Diez, Multivalent electrostatic microtubule interactions of synthetic peptides are sufficient to mimic advanced MAP-like behavior. Mol. Biol. Cell 30, 2953–2968 (2019).

58. E. Prezel, A. Elie, J. Delaroche, V. Stoppin-Mellet, C. Bosc, L. Serre, A. Fourest-Lieuvin, A. Andrieux, M. Vantard, I. Arnal, Tau can switch microtubule network organizations: from random networks to dynamic and stable bundles. Mol. Biol. Cell 29, 154–165 (2018).

59. J. M. Solowska, M. D’Rozario, D. C. Jean, M. W. Davidson, D. R. Marenda, P. W. Baas, Pathogenic mutation of spastin has gain-of-function effects on microtubule dynamics. J. Neurosci. 34, 1856–1867 (2014).

60. R. Ayukawa, S. Iwata, H. Imai, S. Kamimura, M. Hayashi, K. X. Ngo, I. Minoura, S. Uchimura, T. Makino, M. Shirouzu, H. Shigematsu, K. Sekimoto, B. Gigant, E. Muto, GTP-dependent formation of straight tubulin oligomers leads to microtubule nucleation. J. Cell Biol. 220 (2021).

61. D. L. Sackett, B. Bhattacharyya, J. Wolff, Tubulin subunit carboxyl termini determine polymerization efficiency. J. Biol. Chem. 260, 43–45 (1985).

62. C. P. Fees, J. K. Moore, Regulation of microtubule dynamic instability by the carboxy-terminal tail of β-tubulin. Life Sci. Alliance 1 (2018).

63. T.-O. Buchholz, M. Jordan, G. Pigino, F. Jug, “Cryo-CARE: Content-aware image restoration for cryo-transmission electron microscopy data” in 2019 IEEE 16th International Symposium on Biomedical Imaging (ISBI 2019) (IEEE, 2019), pp. 502–506.

64. S. S. Iyer, F. Chen, F. E. Ogunmolu, S. Moradi, V. A. Volkov, E. J. van Grinsven, C. van Hoorn, J. Wu, N. Andrea, S. Hua, K. Jiang, I. Vakonakis, M. Potočnjak, F. Herzog, B. Gigant, N. Gudimchuk, K. E. Stecker, M. Dogterom, M. O. Steinmetz, A. Akhmanova, Centriolar cap proteins CP110 and CPAP control slow elongation of microtubule plus ends. J. Cell Biol. 224 (2025).

65. R. Maan, L. Reese, V. A. Volkov, M. R. King, E. O. van der Sluis, N. Andrea, W. H. Evers, A. J. Jakobi, M. Dogterom, Multivalent interactions facilitate motor-dependent protein accumulation at growing microtubule plus-ends. Nat. Cell Biol. 25, 68–78 (2023).

66. D. J. Odde, L. Cassimeris, H. M. Buettner, Kinetics of microtubule catastrophe assessed by probabilistic analysis. Biophys. J. 69, 796–802 (1995).

67. A. Burt, B. Toader, R. Warshamanage, A. von Kügelgen, E. Pyle, J. Zivanov, D. Kimanius, T. A. M. Bharat, S. H. W. Scheres, An image processing pipeline for electron cryo- tomography in RELION-5. FEBS Open Bio 14, 1788–1804 (2024).

68. Y.-W. Kuo, J. Howard, Cutting, amplifying, and aligning microtubules with severing enzymes. Trends Cell Biol. 31, 50–61 (2021).

69. J. Chen, A. J. Noble, J. Y. Kang, S. A. Darst, Eliminating effects of particle adsorption to the air/water interface in single-particle cryo-electron microscopy: Bacterial RNA polymerase and CHAPSO. J. Struct. Biol. X 1, 100005 (2019).

70. R. Zhang, B. LaFrance, E. Nogales, Separating the effects of nucleotide and EB binding on microtubule structure. Proc. Natl. Acad. Sci. U. S. A. 115, E6191–E6200 (2018).

71. M. Igaev, H. Grubmüller, Bending-torsional elasticity and energetics of the plus-end microtubule tip. Proc. Natl. Acad. Sci. U. S. A. 119, e2115516119 (2022).

72. M. Kalutskii, H. Grubmüller, V. A. Volkov, M. Igaev, Microtubule dynamics are defined by conformations and stability of clustered protofilaments. Proc. Natl. Acad. Sci. U. S. A. 122, e2424263122 (2025).

73. M. Knossow, V. Campanacci, L. A. Khodja, B. Gigant, The mechanism of tubulin assembly into microtubules: Insights from structural studies. iScience 23, 101511 (2020).

74. B. J. LaFrance, J. Roostalu, G. Henkin, B. J. Greber, R. Zhang, D. Normanno, C. O. McCollum, T. Surrey, E. Nogales, Structural transitions in the GTP cap visualized by cryo-electron microscopy of catalytically inactive microtubules. Proc. Natl. Acad. Sci. U. S. A. 119 (2022).

75. B. Gigant, P. A. Curmi, C. Martin-Barbey, E. Charbaut, S. Lachkar, L. Lebeau, S. Siavoshian, A. Sobel, M. Knossow, The 4 Å X-ray structure of a tubulin:Stathmin-like domain complex. Cell 102, 809–816 (2000).

76. E. Eren, N. R. Watts, D. Randazzo, I. Palmer, D. L. Sackett, P. T. Wingfield, Structural basis of microtubule depolymerization by the kinesin-like activity of HIV-1 Rev. Structure 31, 1233–1246.e5 (2023).

77. A. B. Asenjo, C. Chatterjee, D. Tan, V. DePaoli, W. J. Rice, R. Diaz-Avalos, M. Silvestry, H. Sosa, Structural model for tubulin recognition and deformation by kinesin-13 microtubule depolymerases. Cell Rep. 3, 759–768 (2013).

78. G. M. Alushin, G. C. Lander, E. H. Kellogg, R. Zhang, D. Baker, E. Nogales, High-resolution microtubule structures reveal the structural transitions in αβ-tubulin upon GTP hydrolysis. Cell 157, 1117–1129 (2014).

79. S. Bodakuntla, K. Taira, Y. Yamada, P. Alvarez-Brecht, A. K. Cada, N. Basnet, R. Zhang, A. Martinez-Sanchez, C. Biertümpfel, N. Mizuno, In situ structural mechanism of epothilone-B-induced CNS axon regeneration. Nature, 1–11 (2025).

80. E. M. Mandelkow, A. Harmsen, E. Mandelkow, J. Bordas, X-ray kinetic studies of microtubule assembly using synchrotron radiation. Nature 287, 595–599 (1980).

81. J. Aiken, E. L. F. Holzbaur, Spastin locally amplifies microtubule dynamics to pattern the axon for presynaptic cargo delivery. Curr. Biol., doi: 10.1016/j.cub.2024.03.010 (2024).

82. L. I. Binder, A. Frankfurter, L. I. Rebhun, The distribution of tau in the mammalian central nervous system. J. Cell Biol. 101, 1371–1378 (1985).

83. M. Iwata, S. Watanabe, A. Yamane, T. Miyasaka, H. Misonou, Regulatory mechanisms for the axonal localization of tau protein in neurons. Mol. Biol. Cell 30, 2441–2457 (2019).

84. A. Hernández-Vega, M. Braun, L. Scharrel, M. Jahnel, S. Wegmann, B. T. Hyman, S. Alberti, S. Diez, A. A. Hyman, Local nucleation of microtubule bundles through tubulin concentration into a condensed tau phase. Cell Rep. 20, 2304–2312 (2017).

85. V. Siahaan, R. Tan, T. Humhalova, L. Libusova, S. E. Lacey, T. Tan, M. Dacy, K. M. Ori-McKenney, R. J. McKenney, M. Braun, Z. Lansky, Microtubule lattice spacing governs cohesive envelope formation of tau family proteins. Nat. Chem. Biol. 18, 1224–1235 (2022).

86. V. Siahaan, J. Krattenmacher, A. A. Hyman, S. Diez, A. Hernández-Vega, Z. Lansky, M. Braun, Kinetically distinct phases of tau on microtubules regulate kinesin motors and severing enzymes. Nat. Cell Biol. 21, 1086–1092 (2019).

87. E. J. Lawrence, G. Arpag, C. Arnaiz, M. Zanic, SSNA1 stabilizes dynamic microtubules and detects microtubule damage. Elife 10 (2021).

88. U. Goyal, B. Renvoisé, J. Chang, C. Blackstone, Spastin-interacting protein NA14/SSNA1 functions in cytokinesis and axon development. PLoS One 9, e112428 (2014).

89. Y. Zhang, X. He, J. Zou, J. Yang, A. Ma, M. Tan, Phosphorylation mutation impairs the promoting effect of spastin on neurite outgrowth without affecting its microtubule severing ability. Eur. J. Histochem. 67 (2023).

90. L. Chen, H. Wang, S. Cha, J. Li, J. Zhang, J. Wu, G. Guo, J. Zhang, Phosphorylation of Spastin promotes the surface delivery and synaptic function of AMPA receptors. Front. Cell. Neurosci. 16, 809934 (2022).

91. Z.-S. Ji, Q.-L. Liu, J.-F. Zhang, Y.-H. Yang, J. Li, G.-W. Zhang, M.-H. Tan, H.-S. Lin, G.-Q. Guo, SUMOylation of spastin promotes the internalization of GluA1 and regulates dendritic spine morphology by targeting microtubule dynamics. Neurobiol. Dis. 146, 105133 (2020).

92. B. Lacroix, J. van Dijk, N. D. Gold, J. Guizetti, G. Aldrian-Herrada, K. Rogowski, D. W. Gerlich, C. Janke, Tubulin polyglutamylation stimulates spastin-mediated microtubule severing. J. Cell Biol. 189, 945–954 (2010).

93. C. Janke, The tubulin code: molecular components, readout mechanisms, and functions. J. Cell Biol. 206, 461–472 (2014).

94. A. M. Shred, N. E. Vangos, A. N. Bayne, S. C. Tetlalmatzi, W. Peng, J.-F. Trempe, D. Sept, M. A. Cianfrocco, G. J. Brouhard, Conformational flexibility of tubulin dimers regulates the transitions of microtubule dynamic instability, bioRxivorg (2025)p. 2025.06.30.662375.

## References

1. M. Andreu-Carbó, S. Fernandes, C. Aumeier, Two-color in vitro assay to visualize and quantify microtubule shaft dynamics. STAR Protoc. 3, 101320 (2022).

2. A. Burt, B. Toader, R. Warshamanage, A. von Kügelgen, E. Pyle, J. Zivanov, D. Kimanius, T. A. M. Bharat, S. H. W. Scheres, An image processing pipeline for electron cryo-tomography in RELION-5. FEBS Open Bio 14, 1788–1804 (2024).

3. N. Sofroniew, T. Lambert, G. Bokota, J. Nunez-Iglesias, P. Sobolewski, A. Sweet, L. Gaifas, K. Evans, A. Burt, D. Doncila Pop, K. Yamauchi, M. Weber Mendonça, L. Liu, G. Buckley, W.-M. Vierdag, T. Monko, L. Royer, A. Can Solak, K. I. S. Harrington, J. Ahlers, D. Althviz Moré, O. Amsalem, A. Anderson, A. Annex, C. Aronssohn, P. Boone, J. Bragantini, M. Bussonnier, C. Caporal, J. Eglinger, A. Eisenbarth, J. Freeman, C. Gohlke, K. Gunalan, Y. O. Halchenko, H. Har-Gil, M. Harfouche, V. Hilsenstein, K. Hutchings, J. Lauer, G. Lichtner, H. Liu, Z. Liu, A. Lowe, L. Marconato, S. Martin, A. McGovern, L. Migas, N. Miller, S. Miñano, H. Muñoz, J.-H. Müller, C. Nauroth-Kreß, H. A. Obenhaus, D. Palecek, C. Pape, E. Perlman, K. Pevey, G. Peña-Castellanos, A. Pierré, D. Pinto, J. Rodríguez-Guerra, D. Ross, C. T. Russell, J. Ryan, G. Selzer, M. B. Smith, P. Smith, K. Sofiiuk, J. Soltwedel, D. Stansby, J. Vanaret, P. Wadhwa, M. Weigert, C. Willing, J. Windhager, P. Winston, R. Zhao, Napari: A Multi-Dimensional Image Viewer for Python (Zenodo, 2025; 10.5281/zenodo.15779115).

4. E. C. Meng, T. D. Goddard, E. F. Pettersen, G. S. Couch, Z. J. Pearson, J. H. Morris, T. E. Ferrin, UCSF ChimeraX: Tools for structure building and analysis. Protein Sci. 32, e4792 (2023).

5. D. N. Mastronarde, SerialEM: A program for automated tilt series acquisition on Tecnai microscopes using prediction of specimen position. Microsc. Microanal. 9, 1182–1183 (2003).

6. D. Liebschner, P. V. Afonine, M. L. Baker, G. Bunkóczi, V. B. Chen, T. I. Croll, B. Hintze, L. W. Hung, S. Jain, A. J. McCoy, N. W. Moriarty, R. D. Oeffner, B. K. Poon, M. G. Prisant, R. J. Read, J. S. Richardson, D. C. Richardson, M. D. Sammito, O. V. Sobolev, D. H. Stockwell, T. C. Terwilliger, A. G. Urzhumtsev, L. L. Videau, C. J. Williams, P. D. Adams, Macromolecular structure determination using X-rays, neutrons and electrons: recent developments in Phenix. Acta Crystallogr. D Struct. Biol. 75, 861–877 (2019).

7. P. J. A. Cock, T. Antao, J. T. Chang, B. A. Chapman, C. J. Cox, A. Dalke, I. Friedberg, T. Hamelryck, F. Kauff, B. Wilczynski, M. J. L. de Hoon, Biopython: freely available Python tools for computational molecular biology and bioinformatics. Bioinformatics 25, 1422– 1423 (2009).

8. P. Virtanen, R. Gommers, T. E. Oliphant, M. Haberland, T. Reddy, D. Cournapeau, E. Burovski, P. Peterson, W. Weckesser, J. Bright, S. J. van der Walt, M. Brett, J. Wilson, K. J. Millman, N. Mayorov, A. R. J. Nelson, E. Jones, R. Kern, E. Larson, C. J. Carey, İ. Polat, Y. Feng, E. W. Moore, J. VanderPlas, D. Laxalde, J. Perktold, R. Cimrman, I. Henriksen, E. A. Quintero, C. R. Harris, A. M. Archibald, A. H. Ribeiro, F. Pedregosa, P. van Mulbregt, SciPy 1.0 Contributors, SciPy 1.0: fundamental algorithms for scientific computing in Python. Nat. Methods 17, 261–272 (2020).

9. B. J. LaFrance, J. Roostalu, G. Henkin, B. J. Greber, R. Zhang, D. Normanno, C. O. McCollum, T. Surrey, E. Nogales, Structural transitions in the GTP cap visualized by cryo-electron microscopy of catalytically inactive microtubules. Proc. Natl. Acad. Sci. U. S. A. 119 (2022).

